# Glyoxylic acid overcomes 1-MCP-induced blockage of fruit ripening in *Pyrus communis L*. var. ‘D’Anjou’

**DOI:** 10.1101/852954

**Authors:** Seanna Louise Hewitt, Amit Dhingra

## Abstract

1-methylcyclopropene (1-MCP) in an ethylene receptor antagonist which blocks ethylene perception and downstream ripening responses in climacteric fruit imparting a longer shelf life. However, in European pear, application of 1-MCP irreversibly obstructs the onset of system 2 ethylene production resulting in perpetually unripe fruit with undesirable quality. Application of exogenous ethylene, carbon dioxide and treatment to high temperatures is not able to reverse the blockage in ripening. We recently reported that during cold conditioning, activation of AOX occurs pre-climaterically. In this study we report that activation of AOX via exposure of 1-MCP treated ‘D’Anjou’ pear fruit to glyoxylic acid triggers an accelerated ripening response. Ripening is consistently evident in decrease of fruit firmness and onset of S1-S2 ethylene transition. Time course ripening related measurements and transcriptomic analysis were performed to assess the effects of glyoxylic acid-driven stimulation of ripening. Transcriptomic and functional enrichment analyses revealed genes and ontologies implicated in glyoxylic acid mediated ripening, including alternative oxidase, TCA cycle, fatty acid metabolism, amino acid metabolism, organic acid metabolism, and ethylene responsive pathways. These observations implicate the glyoxylate cycle as a metabolic hub linking multiple pathways to stimulate ripening through an alternate mechanism. The results provide information regarding how blockage caused by 1-MCP may be circumvented at the metabolic level, thus opening avenues for consistent ripening in pear and possibly other fruit.

## Introduction

Every year, 1.6 billion tons of food goes to waste. This is about one third of the food that is produced for human consumption [1]. Unpredictable ripening of fruit is one of the main causes of loss after harvest. This is particularly true of climacteric fruits, which are characterized by a spike in ethylene biosynthesis, known as system 2 (S2) ethylene production, and a concomitant burst in respiration at the onset of ripening. The ethylene receptor antagonist 1-methylcyclopropene (1-MCP) is used to impart a longer shelf life by limiting the ethylene perception and activation of downstream ripening responses [2-4].

Uniquely, in European pear fruit (*Pyrus communis*), 1-MCP treatment may irreversibly inhibit endogenous or system 2 ethylene production and the respiratory climacteric [5, 6]. Furthermore, exogenous ethylene application does little to affect the capacity of 1-MCP-treated pears to ripen [7, 8]. This observation suggests that 1-MCP, which has been classically understood only in the context of its identity as an ethylene receptor antagonist [9], seems to exert additional metabolic consequences. This presents a challenge to the U.S. pear industry, as 1-MCP pear fruit fail to ripen properly and do not achieve the desired buttery consistency, aromatics and flavor profile to meet consumer satisfaction standards [10]. Recent research has lent support to the concept of differential effect of 1-MCP on ethylene biosynthetic pathways and signal transduction networks in different climacteric fruits, including peach, pear, apple and tomato [11-13]. Attempts to metabolically reverse the ethylene antagonism of 1-MCP in pear and other climacteric fruit have not been reported previously.

In order to ripen, European pears require a genetically pre-determined amount of cold temperature exposure, known as conditioning [14]. Recent research revealed that AOX1, the key protein in the cyanide resistant alternative respiratory pathway, is elevated significantly in transcript expression during the pre-climacteric stages of cold conditioning in ‘D’Anjou’ and ‘Bartlett’ pear [15]. Climacteric bursts in respiration have been observed in mango and apple fruit and are attributed in part to enhanced post-climacteric AOX activity [16, 17]. It has been proposed that AOX and cytochrome c pathway activity may occur simultaneously in the ripening process, with the alternative respiratory pathway playing a greater role in the senescent processes following the climacteric burst [16, 17].

Based on the recent findings in pear, it appears that AOX may play a role in the pre-climacteric stage in fruits that require conditioning in order to ripen. Chemical genomics approaches targeting AOX for pre-climacteric ripening stimulation identified glyoxylic acid (GLA), the key metabolic intermediate in the glyoxylate cycle, as an activator ripening in 1-MCP treated pear fruit [18]. The glyoxylate cycle is a shortcut in the TCA cycle used by bacteria and plants for gluconeogenesis and carbohydrate synthesis via β-oxidation of fatty acids to Ac-CoA [19-21]. Thus far, the role of the glyoxylate cycle in ripening is largely unexplored, with no studies of metabolic override of 1-MCP ripening blockage reported in present literature.

In this study, the hypothesis that GLA treatment will activate AOX expression and facilitate identification of additional significant genes and gene networks that enable override of 1-MCP blockage of ripening was evaluated. Established indicators of ripening, including fruit firmness, total soluble solids, and internal ethylene production along with RNAseq were analyzed over the course of ripening post-GLA treatment.

## Results and Discussion

Glyoxylic acid treatment resulted in significant decrease in firmness (p<0.05) of 1-MCP treated ‘D’Anjou’ in comparison with the control in each of the three experiments conducted in 2018 (Figure 1). Likewise, internal ethylene peaked significantly (p<0.05) in the GLA-treated pears, a response characteristic of the S1-S2 ethylene transition and the ripening climacteric (Figure 1). °Brix did not change significantly for either the treated or control pears throughout the duration of the experiment (Supplementary File 1). HPLC mean glucose and fructose values were elevated in the 3% GLA treated fruit in comparison with the control despite lack of statistically significant difference between treatment and control groups. Furthermore, analysis of organic acids revealed increased mean malic acid (though not statistically significant) and highly increased citric acid production (p<0.05) in the treated fruit vs the control fruit throughout the ripening time course (Supplementary File 2). Increase in citric acid implicates activation of TCA cycle and respiration upon GLA treatment. Fruit tissues, including peel and flesh, sampled from the 3% GLA treated fruits and from the control fruits were used for a time-course transcriptomic analysis.

**Figure 1.**
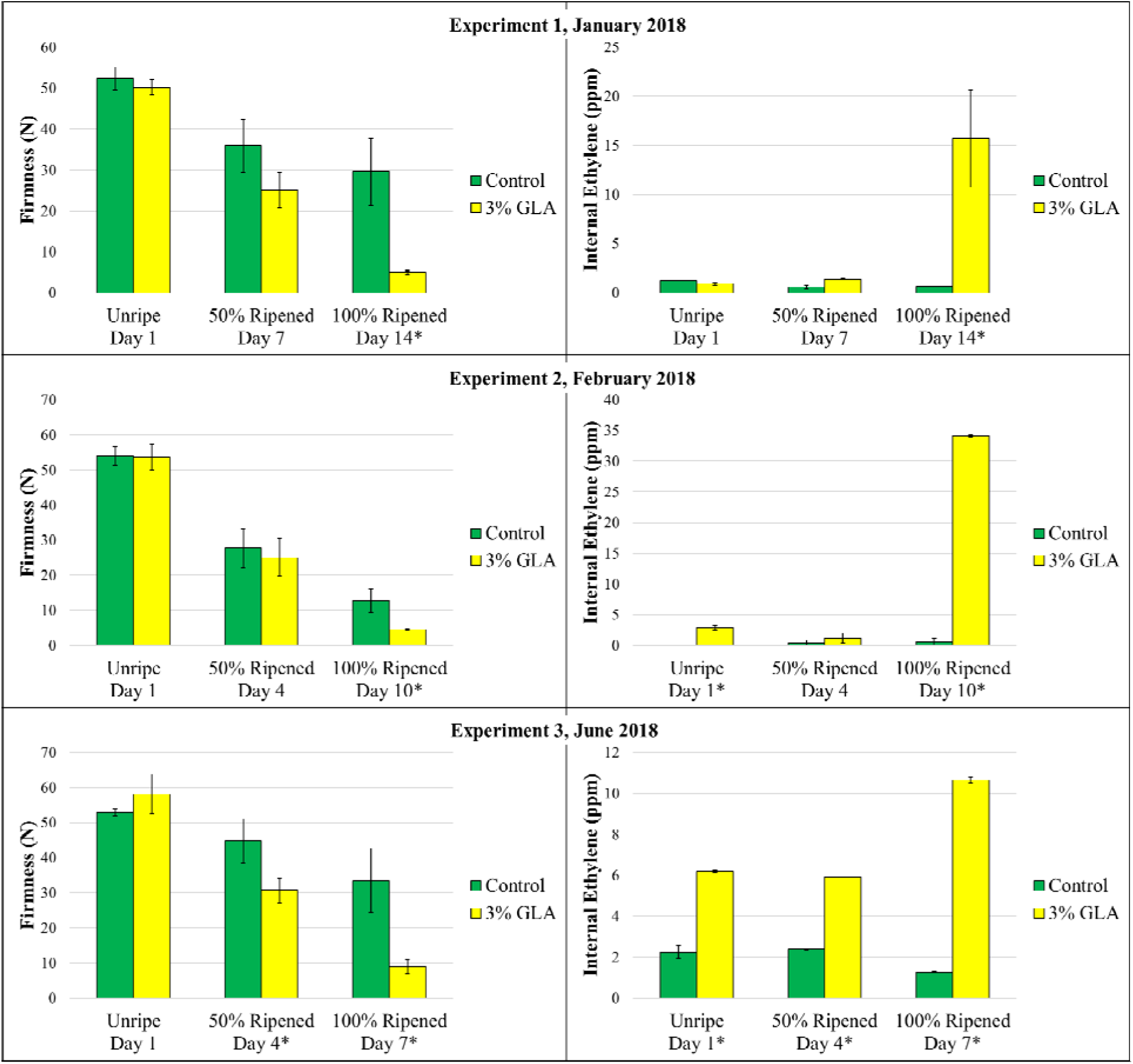
Firmness and internal ethylene of 1-MCP treated ‘D’Anjou pears following treatment with 3% GLA in January 2018 (A), February 2018 (B), and June 2018 (C). Asterisks indicate statistically significant difference at the respective time points (p<0.05).

The duration that each experimental group of pear fruit was held in controlled atmosphere (CA) storage prior to GLA application must also be considered. For delaying ripening of pear fruit, 2-3% oxygen and 0-1% carbon dioxide are the established ranges for CA storage [7, 22], and length of time in CA could impact the fruit’s propensity to ripen. Observations in this study indicated that the overall ripening time course, both in GLA-treated fruit and in control fruit, was shorter the longer the fruit had been in CA storage prior to the start of an experiment. This phenomenon necessitated modified sampling timepoints for each experiment described in the methods section. Treatment endpoint images and analysis of experiments conducted in 2017 that led to optimization of GLA treatment strategy can be found in Supplementary File 3.

### Transcriptomics

RNAseq assembly generated 148,946 contigs (Supplementary File 4). Using the time course differential expression analysis feature in OmicsBox, 5,912 contigs were identified as being differentially expressed over time in the 3% GLA-treated fruit in comparison with the control in all three ripening experiments (Supplementary File 5). To better understand the mechanisms underlying GLA induction of pear fruit ripening, the expression of genes/contigs associated with TCA cycle, glyoxylate cycle, lipid metabolism, AOX and mitochondrial electron transport, sugar/starch breakdown, and organic acid metabolism was analyzed (Table 1). Among the differentially expressed contigs (DECs) were ethylene metabolism-associated contigs and amino acid precursors, numerous ethylene response factors (*ERFs*), a pear *AOX* homolog (ubiquinol oxidase), TCA cycle and glyoxylate cycle-associated contigs, and lipid metabolism-associated contigs. Many of the identified DECs displayed heightened expression in the 3% GLA-treated fruit in comparison with the control fruit at the ‘Unripe’ stage, which was sampled immediately following the 16 hour GLA humidification treatment, indicating that this set of DECs directly responded to the application of GLA, but decreased in expression throughout the time course. Such genes with significantly heightened expression in the GLA-treated fruit immediately following treatment included: *AOX1*, several *ERF* transcripts, *RAN1*, isocitrate dehydrogenase, lipoxygenase, and *AOS1*. DECs which displayed delayed transcriptional responses, which became evident at the ‘50% Ripened’ to ‘100% Ripened’ included: ethylene biosynthetic enzymes, ACS and ACO, and citrate synthase, which displayed heightened expression in the treatment group in later stages of ripening.

**Table 1.** Summary of differentially expressed contigs discussed in the manuscript. Information includes associated pathway, full and abbreviated names, contig number (corresponding to sequences, annotations, and expression values in Supplementary File 5), length, and indication of significant trends.

| Associated pathway | Gene/Contig Name | Abbreviation | Contig # | Contig Length (bp) | Significant Trend<br>3% Glyoxylic Acid<br>(R>0.8) |
| --- | --- | --- | --- | --- | --- |
| Alternative respiration | Ubiquinol oxidase, mitochondrial-like | AOX1 | 22331 | 1156 |  |
| Aspartate, cysteine, methionine metabolism | Aspartate aminotransferase, cytoplasmic | cytAAT | 18640 | 503 | Linear, Quadratic |
| Aspartate, cysteine, methionine metabolism | Bifunctional aspartate aminotransferase | BiAAT | 10407 | 850 |  |
| Aspartate, cysteine, methionine metabolism | SAM synthase 2 | SAM synthase 2 | 2250 | 1734 |  |
| Aspartate, cysteine, methionine metabolism | Probable pyridoxal 5'-phosphate synthase subunit PDX2 | PDX2 | 6815 | 1473 |  |
| Aspartate, cysteine, methionine metabolism | Cystathionine gamma-synthase 1, chloroplastic | CγS | 9947 | 1598 | |
| Ethylene biosynthesis | 1-aminocyclopropane-1-carboxylate synthase | ACS | 61428 | 1380 | Linear |
| Ethylene biosynthesis | ACC oxidase | ACO | 3326 | 595 |  |
| Ethylene perception | Copper-transporting ATPase RAN1 | RAN1 | 20950 | 2399 | Linear |
| Ethylene signaling/response | Ethylene-responsive transcription factor ERF073-like | ERF073-like | 43790 | 243 |  |
| Ethylene signaling/response | ETHYLENE INSENSITIVE 3-like 1 protein | EIN3-like 1 | 3680 | 454 | Linear, Quadratic |
| Ethylene signaling/response | Ethylene-responsive transcription factor 3-like | ERF 3-like | 11186 | 1079 |  |
| Ethylene signaling/response | Ethylene-responsive transcription factor 4 isoform X1 | ERF4 isoform x1 | 20105 | 297 |  |
| Ethylene signaling/response | Ethylene-responsive transcription factor 5-like | ERF5-like | 54832 | 1214 | Linear |
| Ethylene signaling/response | Ethylene-responsive transcription factor ABR1-like | ERF ABR1-like | 7129 | 1416 |  |
| Ethylene signaling/response | Ethylene-responsive transcription factor ERF096-like | ERF096-like | 35082 | 706 |  |
| Ethylene signaling/response | Serine/threonine-protein kinase CTR1 isoform X1 | CTR1 isoform x1 | 20857 | 650 |  |
| Fatty Acid/Oxylipin metabolism | Lipoxygenase | LOX1 | 4033 | 2690 |  |
| Fatty Acid/Oxylipin metabolism | Allene oxide synthase | AOS | 3632 | 1639 |  |
| Fatty Acid/Oxylipin metabolism | Allene oxide cyclase 4, chloroplastic | AOC4 | 2502 | 979 | Quadratic |
| Fatty Acid/Oxylipin metabolism | Probable linoleate 9S-lipoxygenase 5 | Linoleate 9S-LOX 5 | 15488 | 3275 | Linear |
| Fatty Acid/Oxylipin metabolism | Linoleate 13S-lipoxygenase 3-1, chloroplastic-like | Linoleate 13S-LOX 3-1 | 10268 | 2980 |  |
| Glyoxylate cycle | Citrate synthase, glyoxysomal | gCS | 3891 | 1784 |  |
| Glycolysis/Gluconeogenesis | Phosphoenolpyruvate carboxylase kinase 1-like | PEPCK1-like | 10747 | 754 |  |
| Glycolysis/Gluconeogenesis | ATP-dependent 6-phosphofructokinase 3-like | PFK3-like | 19400 | 904 |  |
| Glycolysis/Gluconeogenesis | Fructose-1,6-bisphosphatase, cytosolic | F-1,6-Bpase | 42519 | 255 | Linear, Quadratic |
| TCA cycle | Fumarate hydratase 1, mitochondrial | mFH1 | 6653 | 845 | Linear |
| TCA cycle | Isocitrate dehydrogenase [NADP] | IDH | 7737 | 2131 |  |
| TCA/Glyoxylate cycle | Citrate synthase, mitochondrial | mCS | 1515 | 418 |  |
| TCA/Glyoxylate cycle | Malate dehydrogenase | MDH | 9623 | 2234 | Linear |

### Glyoxylic acid activation of AOX

Alterative oxidase (AOX) has been implicated previously in ethylene biosynthetic processes, the respiratory climacteric and ripening. Immediately following treatment with GLA, *AOX* transcript abundance was higher, with its expression decreasing over time in both treatment and control 1-MCP fruit but remaining consistently higher in the GLA treatment group (Figure 2). This expression trend is reminiscent of the pre-climacteric maxima of AOX transcription in non-1-MCP treated pear fruit that had undergone full conditioning [15, 23]. AOX pathway activity has been shown to contribute to the respiratory climacteric during ripening in tomato, papaya, and mango [16, 24-26]. While GLA activated *AOX*, the results of this transcriptomic analysis suggest that AOX activity in 1-MCP treated pear fruit is potentially regulated in a feed-forward manner as a result of carbon metabolism upstream [24, 25, 27]. This alternative respiratory stimulation may partially account for the enhanced respiratory response and ripening of 1-MCP treated pear [18].

**Figure 2.**
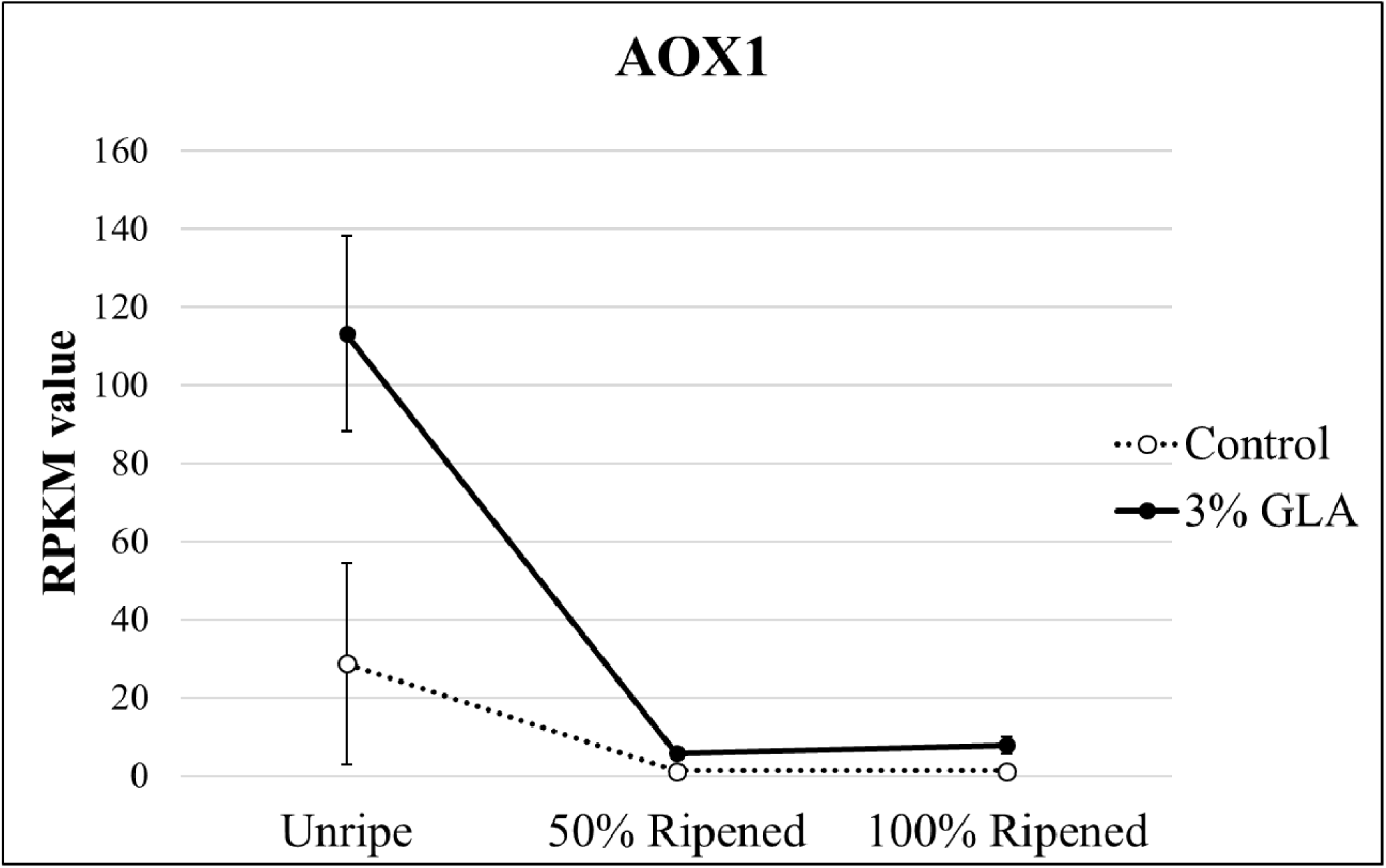
Expression of AOX1 homologue. AOX was significantly differentially expressed over time in the 3% GLA-treated fruit than in the control fruit (p<0.05).

### Gluconeogenesis and glycolytic flux balance

The glyoxylate cycle is directly linked to the production of glucose via gluconeogenesis in certain biological contexts such as fatty acid to sugar conversion during seedling germination [19]. Previously, study of gluconeogenic enzymes in tomato and peach revealed that *PEPCK* abundance increases during ripening of these fruit [28]. It has also been demonstrated that malate and citrate accumulate throughout fruit development and may be utilized both as substrates for respiration in the TCA cycle, as well as for gluconeogenesis during fruit ripening; however, 1-MCP slows the rate of organic acid to sugar conversion via gluconeogenesis which thereby delays ripening [29-31]. Thus, it was of interest to examine expression of genes in the gluconeogenic pathway in addition to the glycolytic pathway.

The control 1-MCP fruit in this experiment displayed stable mean expression levels of *PEPCK* over the time course. In GLA treated 1-MCP fruit, mean expression levels of *PEPCK* were higher than in the control immediately following treatment, and gradually decreased over the time course, ending with very low expression levels in comparison with the control. *Fructose 1,6 bisphosphatase* (*FBPase*), which encodes an enzyme that plays an additional key role in gluconeogenic pathway, displayed elevated expression levels in the control versus the treatment throughout the duration of the experimental time course (Figure 3A). Conversely, *ATP-dependent 6-phosphofructokinase 3-like* (*PFK-like*) DEC exhibited dramatically elevated expression in the GLA-treated fruit immediately following treatment in comparison with the control fruit and decreased over time. PFK catalyzes the first committed step of glycolysis, which leads to the formation of ATP and pyruvate (Figure 3A). The ATP generated through this process can catalyze other ripening related processes.

**Figure 3.**
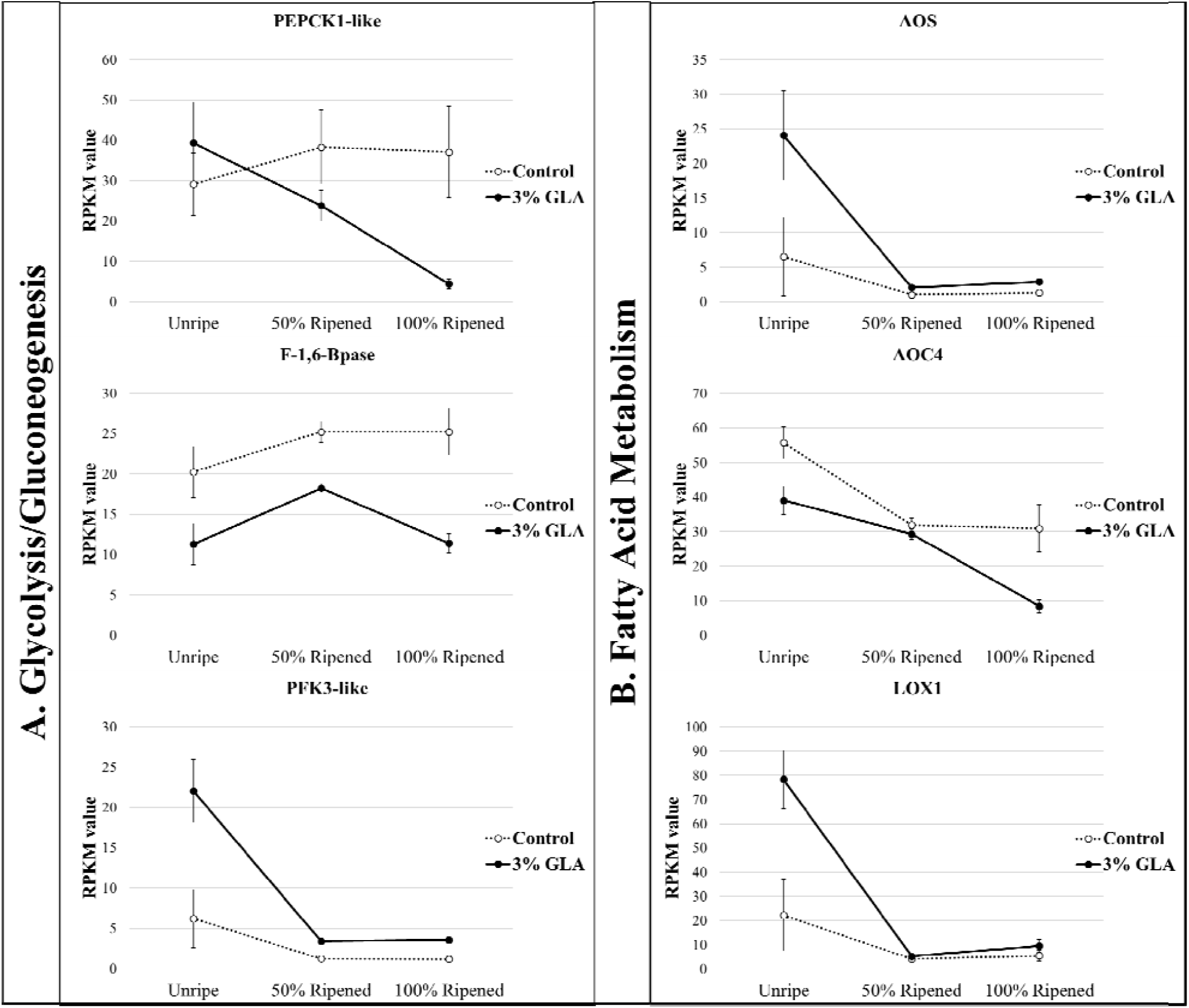
Genes encoding rate limiting gluconeogenic and glycolytic enzymes (column A) and fatty acid and oxylipin metabolic enzymes (column B) that were significantly differentially expressed over time in the 3% GLA-treated fruit versus the control fruit (p<0.05).

Taken together, these results suggest that in GLA-treated 1-MCP pear fruit, flux through the gluconeogenic pathway results in additional allocation of carbon-based metabolites to the TCA cycle, potentially leading to overreduction of the mitochondrial electron transport chain, thereby necessitating increased AOX activity.

### Fatty Acid Metabolism and Oxylipins

The glyoxylate cycle has an important role in β-oxidation of fatty acid under certain biological conditions, seedling germination being one of them, in which production of sugar from other organic substrates is of developmental importance [19, 32]. Addition of GLA could potentially induce multiple metabolic pathways, thereby facilitating a link between fatty acid metabolism and GLA-induced ripening.

Fatty acid oxidation capacity typically is at its highest in pre-climacteric fruit and in fruit entering the climacteric stage [33]. Furthermore, a shift from mitochondrial oxidation of fatty acids to TCA cycle intermediates occurs in the climacteric transition (S1-S2 ethylene biosynthesis) [33]. In mango, addition of glyoxylate stimulated oxidation of fatty acids occurs in a concentration dependent manner [33]; it appears the same may be occurring in GLA treated 1-MCP pear fruit.

Oxidation of unsaturated fatty acids results in the generation of a class of lipophilic signaling molecules called oxylipins. Initial synthesis of these molecules occurs via conversion of polyunsaturated fatty acids to fatty acid hyperperoxides via lipoxygenase (LOX) [34]. The phytohormone jasmonic acid (JA) is among the most well characterized oxylipins, as jasmonates, together with ethylene, are believed to play a role in the early stages of climacteric ripening [35, 36]. Methyl jasmonate increases postharvest shelf life in Chilean strawberry (*Fragaria chiloensis*) by modulating the soluble solid to TA ratios as validated by assessments of quality chemical attributes and decay incidence [37]. JA has been shown to negatively regulate ripening in ‘Bartlett’ pears and is suggested to do so by working either independently or upstream of ethylene biosynthesis [38]. The first specific enzyme, and rate limiting point in JA biosynthesis is allene oxide synthase (AOS) [39]. Expression of AOS was significantly elevated in the GLA-treated fruit in comparison with the control 1-MCP fruit immediately following treatment, suggesting an increased flux through the JA biosynthesis pathway, which could potentially aid in further stimulation of an ethylene response in the treated fruit (Figure 3B). In addition to AOS, several lipoxygenases were also significantly elevated in expression in the treatment versus the control fruit (some at the beginning and others at the end of the time course), indicating the commitment of fatty acid breakdown products to oxylipin metabolism (Figure 3B). Based on these observations, it appears that fatty acid metabolism contributes both to a hormonal response and to increased pools of carbon-metabolites available for TCA cycle, thereby contributing to the multifaceted ripening response stimulated by GLA.

### TCA and glyoxylate cycle metabolism

The glyoxylate cycle is primarily compartmentalized in the peroxisome, however, two of the five enzymes that comprise the cycle are cytosolic (gCS, gICL, gMS, ctAC, ctMDH). Malate is an intermediate in both glyoxylate and mitochondrial TCA cycles and has been suggested to play a role as an important regulatory metabolite during ripening [40, 41]. This important organic acid is involved in many other processes besides the glyoxylate cycle and respiratory metabolism, and therefore, membrane transport is required to shuttle this substrate into other subcellular compartments (Pracharoenwattana & Smith, 2008). Glyoxylate cycle enzyme encoding malate synthase gene expression has been detected specifically in ripening tissue, but this expression varied by ripening stage and has been suggested that expression changes could be stimulated by exogenous ethylene [42, 43]. Alteration in the supply of malate has been shown to affect postharvest storage quality [41]. Like malate, citrate is an intermediate in both TCA and glyoxylate cycles that accumulates in the pre-climacteric stages and is metabolized during ripening and is one of the most prevalent metabolites contributing to flavor and titratable acidity during fruit development [33]. Contigs corresponding to glyoxysomal and mitochondrial citrate synthase enzymes differed significantly in expression over time in GLA-treated 1-MCP fruit versus the control 1-MCP fruit, with increased expression of both observed in the treatment. Additional contigs corresponding to TCA cycle enzymes fumarate hydratase, isocitrate dehydrogenase were also elevated in expression in GLA-treated fruit (Figure 4). It implicated, based on the expression of TCA/glyoxylate cycle genes that GLA could directly affect the flux through these pathways.

**Figure 4.**
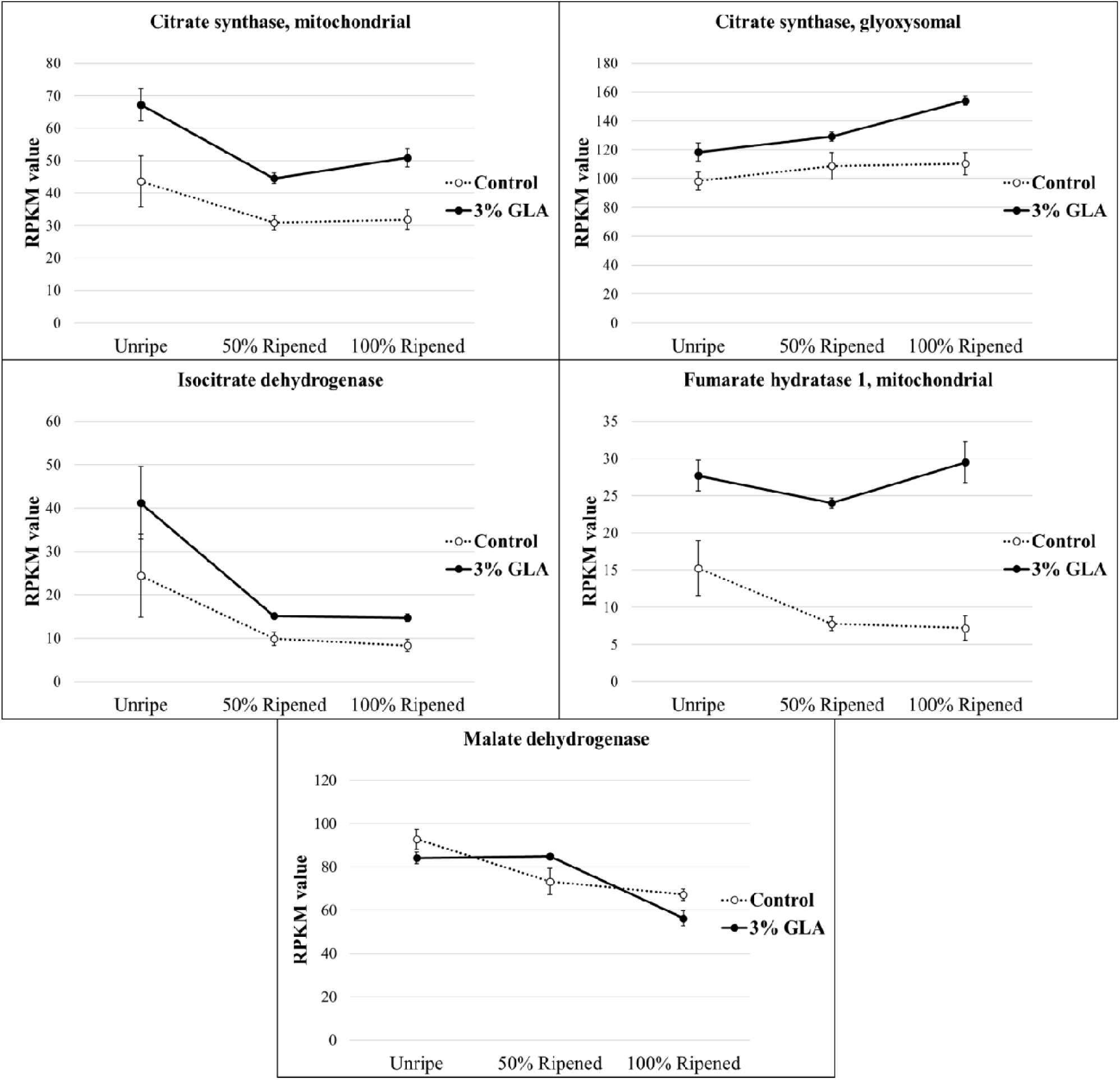
Genes encoding TCA/glyoxylate cycle enzymes that were significantly differentially expressed over time in the 3% GLA-treated fruit versus the control fruit (p<0.05).

### Glyoxylic acid link to ethylene biosynthesis

Methionine cycling, and therefore ethylene production is a process that operates concomitantly with respiration and is co-dependent upon the TCA/glyoxylate cycle compound, oxaloacetate. In addition to serving as a substrate for the formation of citrate, OAA may be converted into intermediates for other metabolic processes [44]. The link between the glyoxylate and TCA cycle and ethylene begin with the conversion of OAA to aspartate via aspartate aminotransferase (AAT) or by bifunctional aspartate/prephenate aminotransferase (PT-AAT/PAT). Aspartate is then converted to homoserine, cystathionine, homocysteine, and then to methionine. Methionine is converted to S-adenosyl-L methionine catalyzed by SAM synthetase, forming the precursor to ACC and ethylene biosynthesis. The enzymes involved in these conversions depend on the co-factor pyridoxal-5’-phosphate (PLP), the production of which is catalyzed by pyridoxine synthases (PDX1 and PDX2) [45, 46]. The heightened expression of genes in the path from OAA to the production of ACC in ethylene biosynthesis in the GLA-treated fruit in comparison with the control 1-MCP fruit suggest that the ripening compound elicits effects that upregulate this pathway via a yet uncharacterized mechanism. *AAT* expression was similar in the treatment and control at the experimental start and end points; however, its expression was significantly higher at the 50% Ripened stage (Supplementary File 6). *AAT/PAT* expression was highest in the GLA-treated fruit immediately following treatment, decreasing throughout the time course, but all the while remaining expressed at higher levels than the control fruit. *CγS* expression remained highest in the GLA treatment over time than the control fruit. The expression pattern of *AAT/PAT, CγS*, and *SAM* as well as the cofactor producing *PDX2*, followed a nearly identical pattern, with highest expression immediately following treatment and decreasing expression throughout the time course (Supplementary File 6). This methionine may feed directly into the methionine cycle, thereby resulting in increased ethylene production when OAA is produced abundantly. The observed changes in expression patterns suggest that increased flux through the glyoxylate/TCA cycles may lead to increased production of OAA, some of which is shunted into the methionine biosynthesis pathway and can be consequently converted to ethylene. Methionine cycling is an ATP-dependent process and in situations where respiration is interrupted, as is in the case of 1-MCP treatment, this fundamental process cannot occur. Activation of the glyoxylate cycle in fruit subjected to respiratory inhibition is the potential source for the ATP necessary for continued methionine cycling and, therefore, ethylene biosynthesis.

Significant changes in the expression of DECs associated with ethylene biosynthesis, perception, and signaling were identified throughout the ripening time course following GLA treatment, including those corresponding to ethylene biosynthetic genes *ACO1* and *ACS1*, ethylene perception-associated *RAN1* and *CTR1* (Figure 5), and numerous ethylene response factors (Supplementary File 7).

**Figure 5.**
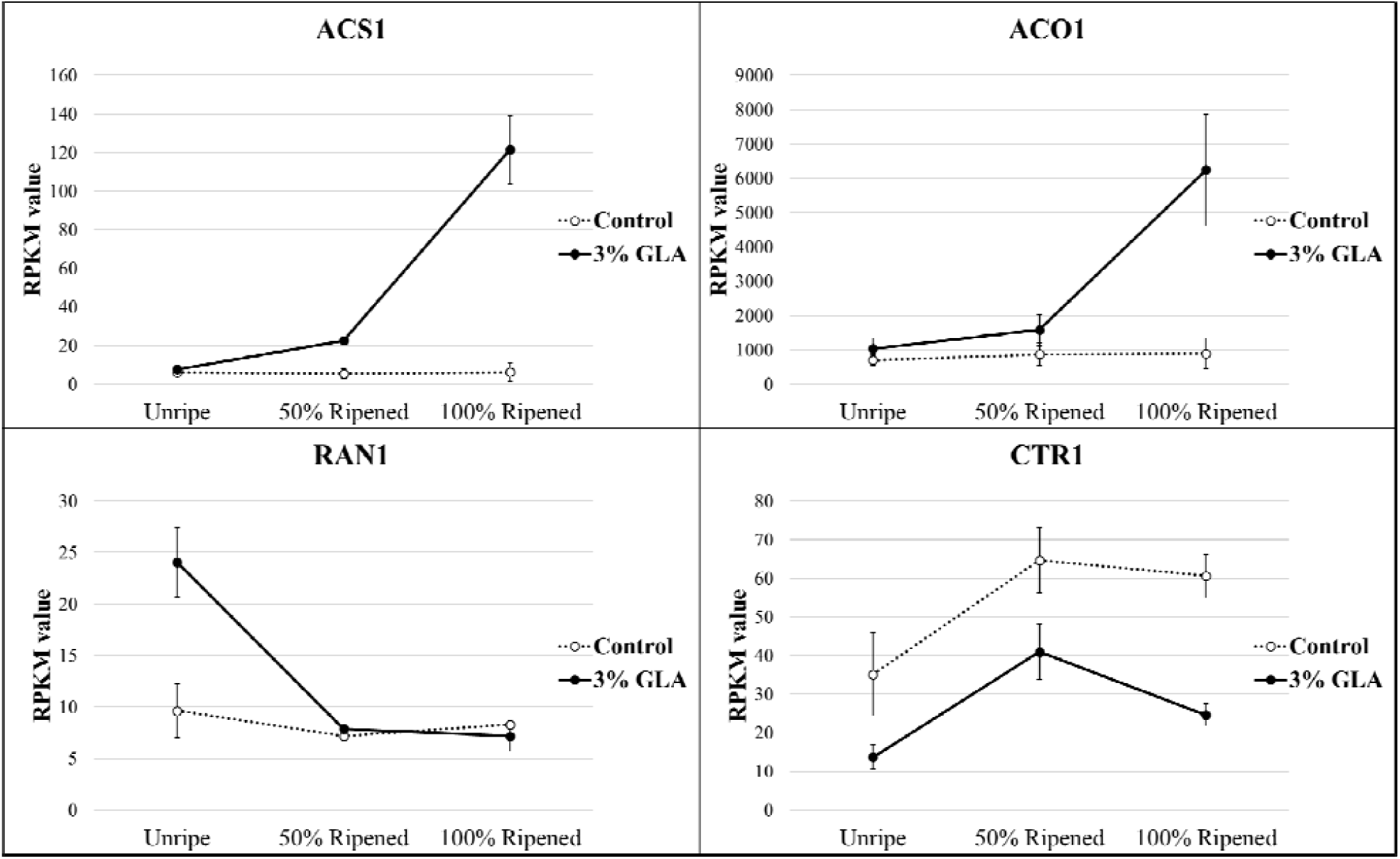
Expression of genes involved in ethylene biosynthesis and perception that were significantly differentially expressed over time in the treatment group versus the control (p<0.05).

### Functional Enrichment Analysis

GO enrichment analysis was conducted using the differentially expressed contigs in the 3% GLA treatment group versus the control group as the test set and the annotated master assembly as the reference set, and the resulting enriched ontologies were reduced to most specific (FDR<0.05). The results provide complementary information to the differential expression analysis, with many of the over and underrepresented terms corresponding to biological processes and molecular functions associated with known and novel ripening related pathways (Supplementary File 8).

Overrepresented ontologies in the 3% GLA-treated fruit included: ‘fruit ripening’ and ‘response to chemical’. These are expected terms, considering that physiological changes indicative of ripening in response to chemical treatment were observed in the treated fruit. Additional overrepresented GO terms included those associated with phytohormone metabolism (‘1-aminocyclopropane-1-carboxylate biosynthetic process’, ‘hormone metabolic process’, and ‘regulation of hormone levels’), ‘aspartate family amino acid biosynthetic process’, metabolism of sulfur compounds (‘sulfate transport, ‘sulfur compound metabolic process’), organic acid metabolism (‘carboxylic acid catabolic process’, ‘oxoacid metabolic process’), lipid metabolism (‘lipid metabolic process’, ‘oxylipin biosynthetic process’). These findings coincide with the results of the differential expression analysis where contigs associated with ethylene biosynthesis and signaling, organic acid metabolism, and lipid metabolism displayed significant differential expression (Figures 3-5), providing support for crosstalk between these seemingly independent pathways during GLA-mediated ripening. Furthermore, results of this analysis again implicate AOX as an important factor in ripening. Sulfur metabolism-associated terms were overrepresented, and sulfur has previously been implicated in the pre-climacteric activation of the AOX respiratory pathway and ripening in cold-conditioned pear fruit [18]. Interestingly, ‘response to cold’, ‘cellular oxidant detoxification’ and ‘antioxidant activity’ were also enriched. While these experiments were conducted at ambient temperature, cold conditioning is required for pear fruit to ripen [5], and recent studies of cold-induced ripening of pear implicate the involvement of AOX for crosstalk with ethylene and scavenging of reactive oxygen species (ROS), thereby alleviating oxidative damage under external stress conditions [15, 47]. Enrichment of these ontologies may indicate that similar mechanisms are activated in both cold and GLA-induced ripening, with both instances involving AOX activity. It raises an interesting possibility that GLA might be able to, in part, replace the requirement of cold conditioning in pear.

Underrepresented GO terms include those associated with maintenance of DNA and chromosomal integrity and organization (‘DNA-dependent DNA replication’, ‘DNA repair’, ‘DNA conformation change’, ‘chromosome modification’, ‘chromatin organization’, and ‘histone modification’) (Supplementary File 8). The decreased representation of these functions in the GLA-treated test set, which exhibited physiological and transcriptomic indications of more rapid senescence, indicates that compositional organization decreases, and cellular entropy increases as ripening progresses. These processes are directly correlated to cell wall softening, and other processes of decompartmentalization in the ripening fruit. Backtracking from the GO enrichment data to corresponding differentially expressed genes may provide additional targets for ripening regulation in a number of related pathways.

### Glyoxylic acid as a metabolic hub

The findings of this study suggest that the propensity of GLA to stimulate a ripening response in pear may result from the centrality of the glyoxylate metabolism to several critical biochemical pathways, as evidenced by the elevation in expression of genes encoding critical rate limiting enzymes in glycolysis, fatty acid metabolism, aspartate/cysteine/methionine metabolism, TCA cycle, and the AOX respiratory pathway immediately following GLA treatment (Figure 6). Interestingly, while such closely associated processes with the glyoxylate cycle appear to be stimulated upon GLA administration, differential expression of the unique glyoxylate cycle enzymes malate synthase and isocitrate lyase was not observed. This finding necessitates further protein expression-based work but could also indicate that following application, GLA is being directly converted into other substrates that can be used for respiration and ethylene biosynthesis—testing the effects of continual administration of GLA at low levels throughout a designated ripening period could shed light on this hypothesis. Such an approach has already been successfully employed with other compounds, including 1-MCP [6]. Additionally, investigating the potential stimulatory effects of exogenously applied TCA and glyoxylate cycle intermediates would add further insight to the understanding of the way in which these pathways and their metabolic intermediates affect critical developmental processes like ripening.

**Figure 6.**
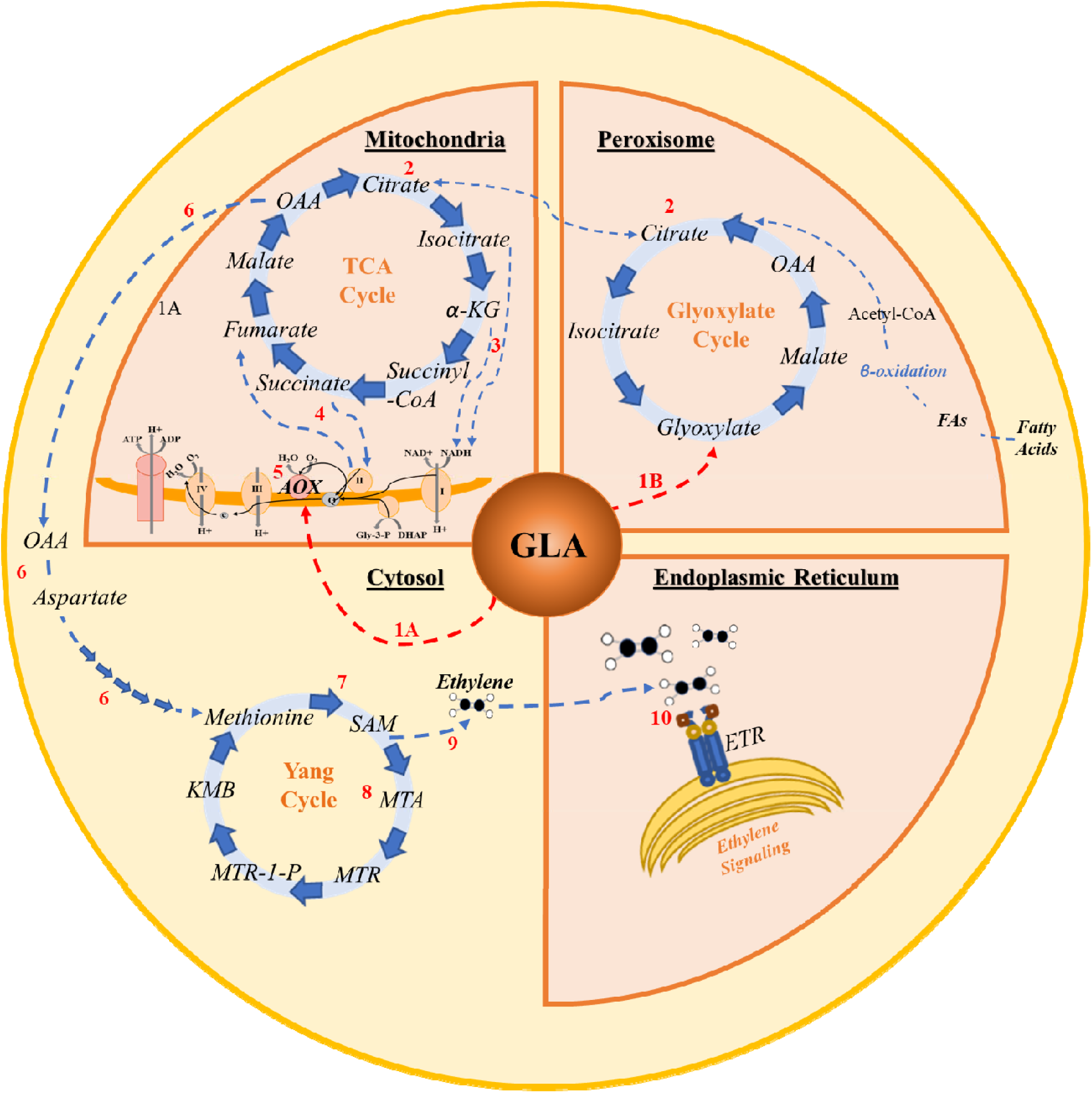
Proposed model for glyoxylic acid as a metabolic hub connecting ripening associated pathways in the peroxisome (glyoxylate cycle and fatty acid β-oxidation), mitochondria (TCA cycle and mETC), cytoplasm (methionine cycle and ethylene biosynthesis), and endoplasmic reticulum (ethylene perception and signaling). (1) Putative sites for direct activation of AOX are indicated by the red dashed arrows. 1A-Glyoxylate has previously been shown to directly activate AOX activity in Arabidopsis [48], and GLA is hypothesized to elicit similar AOX activation in other systems, indicated by a red dashed arrow. 1B-Addition of GLA increases pools of metabolic intermediates in the glyoxylate cycle, resulting in increased flux through the cycle. (2) Production of malate and its subsequent conversion to OAA, perpetuates th glyoxylate cycle, producing intermediates shared by the TCA cycle, including those which can be shuttled between intracellular compartments (e.g. citrate). Thus, GLA leads to increased flux through both glyoxylate and TCA cycles. (3) The oxidative decarboxylations of isocitrate and α-ketoglutarate in the TCA cycle result in the transfer of high-energy electrons to NADH, which is transported to the mitochondrial cytochrome c (CYTc) pathway. (4) Conversion of succinate to fumarate in the TCA cycle results in transfer of additional high energy electrons to complex II of the CYTc pathway. (5) The resulting increased flux of electrons necessitates activation of AOX to prevent overreduction of the CYTc pathway. (6) In addition to respiratory activation, the TCA and glyoxylate cycles share the common intermediate OAA, which can be converted to aspartate and subsequently to the precursors of methionine. (7) Methionine, a precursor for ethylene biosynthesis is converted to SAM. From SAM, methionine is either (8) recycled via the Yang cycle, or (9) converted directly into ethylene. (10) Ethylene binds to ER bound receptors (ETRs), initiating the ethylene signal cascade and downstream ripening-associated responses.

## Conclusion

This study allowed for the identification of interacting networks of genes and pathways that are involved in GLA-mediated activation of AOX and the metabolic override of ripening blockage in pear caused by 1-MCP. Based on prior knowledge of the glyoxylate cycle, its intermediates, and its regulation, it is hypothesized that its mode of action in ripening induction occurs via one or more of the following processes: direct activation of AOX, induction of fatty acid oxidation, provision of organic acid as respiratory substrate, indirect TCA cycle stimulation, or induction of gluconeogenesis [32, 33, 48-51] (Figure 6). Results of time course differential expression analysis of GLA-treated ‘D’Anjou’ 1-MCP pear fruit compared to control 1-MCP fruit lend support to this hypothesis and have provided additional molecular targets for fine-tuned ripening regulation of European pear and other climacteric fruits subjected to 1-MCP treatment.

Ripening in climacteric fruit has primarily been attributed to biosynthesis of ethylene. It is clear, however, that the nuances of ripening extend far beyond ethylene production and respiration [52]. Due to complex requirements for S1-S2 transition in European pear, the fruit may serve as a useful system in which to explore the vagaries of the ripening process in climacteric and non-climacteric fruit [52]. This study addresses a knowledge gap in the ripening process in pear, epitomized by the inability of 1-MCP treated fruit to regain ripening capacity naturally and paralleled with the discovery that GLA can overcome the 1-MCP blockage. Exploring the potential role of glyoxylate as a metabolic hub, its link to ethylene biosynthesis, fatty acid metabolism, glycolysis, and activation of the AOX respiratory pathway in the context of 1-MCP inhibited ripening, provides further insight into mechanism of ripening in European pear and other fruit. It also enriches understanding regarding how ripening control may be better achieved through 1-MCP inhibition and subsequent stimulation of ripening using natural metabolic intermediates.

The role of GLA in stimulation of ripening and the potential genes and pathways implicated in this process provide additional molecular targets for fine-tuned regulation of ripening, and contribute to a body of work in which genetic markers associated with sugars, acids, volatiles, and ripening in general are being mapped [53, 54]. As GLA is a natural plant metabolite and has previously been used as commercial food preservative [15, 55], it is feasible to introduce it as a ripening tool in the tree fruit industry. Translation of this ripening tool set—1-MCP inhibition and GLA activation—to other crops in the future, is expected to provide the opportunity to inhibit ripening at harvest, store fruit for the desired duration or ship it to domestic or international locations. Ripening can be reactivated in a planned and predictable manner, thereby improving consumer satisfaction and ultimately improving postharvest sustainability by reducing the massive waste associated with unpredictable ripening of fruit.

## Materials and Methods

### Acquisition of ‘D’Anjou’ pear fruit

Mature, late season ‘D’Anjou’ pears were obtained from Blue Star Growers (Cashmere, WA) in the Winter/Spring of 2018. Prior to acquisition, pears had been treated with 130ppm 1-MCP and retained in controlled atmosphere storage at 5°C. Following acquisition, pears were transferred to 1°C of the WSU Johnson hall postharvest facility’s controlled atmosphere room for approximately one week prior to initiation of experiments.

### Ultrasonic humidification of ripening compound treatment solutions

Pears were transferred to non-airtight plexiglass tanks (dimensions 40cm × 58cm × 66cm) fitted with an inlet for a tube connected to a Crane Ultrasonic humidifier (Crane USA). The bottom of each humidifier was lined with baker’s drying racks to allow for excess treatment liquid to drip to the bottom of the chamber during the humidification period. Humidifiers were loaded with treatment and control solutions of 3% GLA and deionized water, respectively (Supplementary File 9). Depending on the number of samples to be taken throughout the duration of the experiment, between 32 and 64 pears were treated in each chamber. The pears were treated for a total of 16 hours overnight with their designated ripening compound or control solution.

### Tissue sampling

It was necessary to classify sampling points to be used for sequencing based on percent ripeness. This method of sample point determination has been previously employed in ripening studies [15, 56]. Fruit was classified as ‘Unripe’ on Day 1, immediately upon removal from GLA or control humidification chambers, when firmness between 11-13 lbf (48.9-57.8 N). The classification as ‘50% Ripened’ was given to fruit at the sample point where average firmness of 3% GLA-treated fruit was between 5-7 lbf (22.2-31.1 N). The fruit were classified as ‘100% Ripened’ when the firmness of the 3% GLA-treated fruit were between 1-3 lbf (4.4-13.3N) (Figure 7). Sample analysis stages corresponding to 0, 50, and 100% Ripened for Experiment 1 (January-February 2018) were determined to be Day 1, Day 7 and Day 14; Experiment 2 (February 2018) Day 1, Day 4, and Day 10; and Experiment 3 (June 2019) Day 1, Day 4 and Day 7.

**Figure 7.**
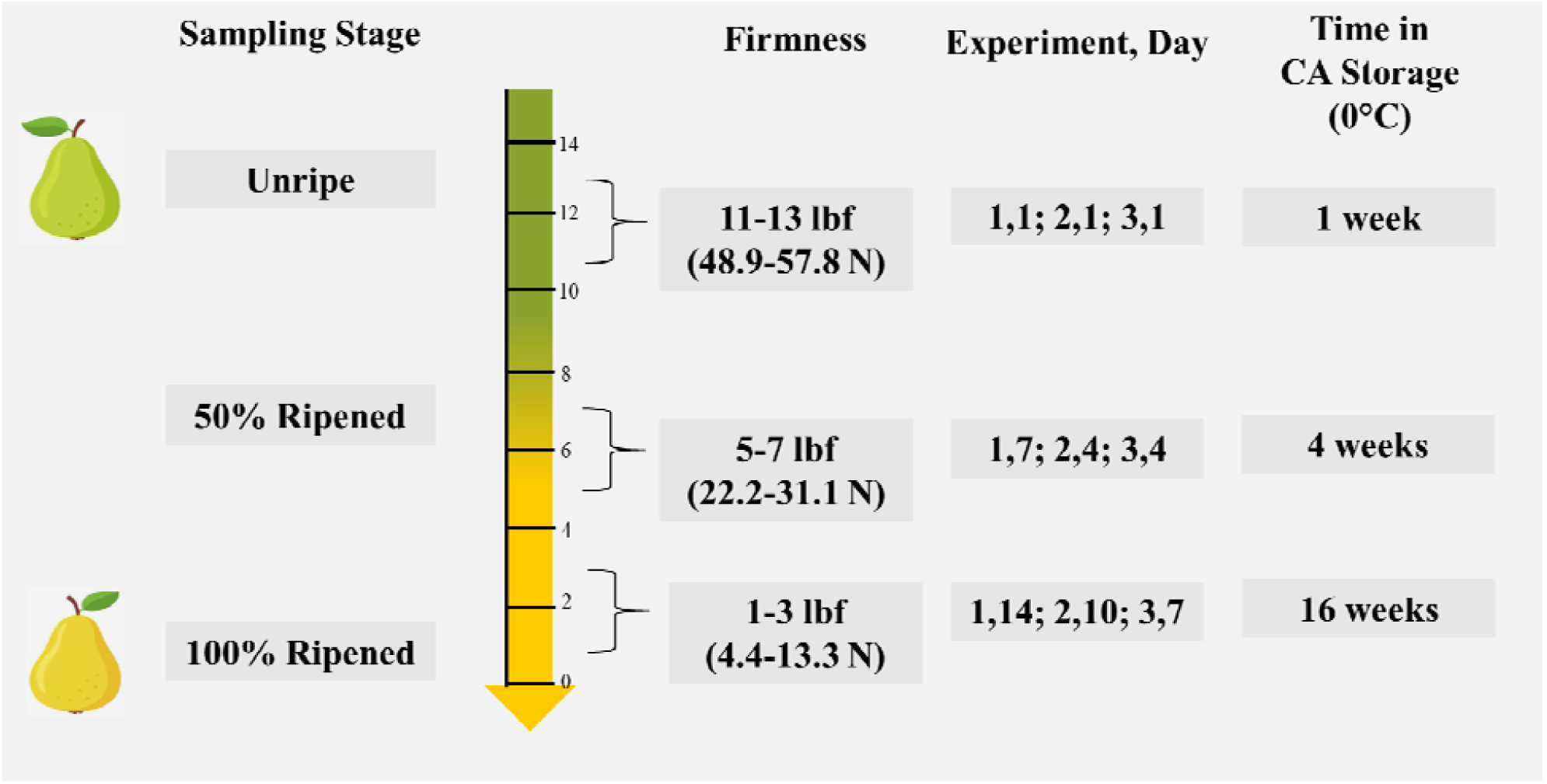
Sampling time-course and experimental replicate specific information for ‘D’Anjou’ pear fruit used in each GLA experiment.

In each experiment, peel tissue was sampled from a 1 cm wide equatorial region of each of 4 fruit per treatment and time point, pooled, flash frozen in liquid nitrogen, and ground using a SPEX Freezer/Mill 6870 (Metuchen, NJ USA). Tissues were stored at -80°C prior to RNA extraction and HPLC analysis.

### Monitoring of carbon dioxide evolution

Following 16-hour humidification treatment, GLA-treated and control pears were weighed and sealed into continuous air flow chambers (four replicates of 4 fruit per replicate), which sample CO_2_ at 4-hour intervals. Fruit respiration rates were determined by calculating mean CO_2_ evolution per kilogram of fruit at each sample time using the previously described methods and sampling system [57].

### Internal ethylene gas chromatographic measurements

Ethylene gas was vacuum extracted from the inside of the fruit using a vacuum aspirator according to previously described methods [58]. One quarter of four separate pear fruit per treatment was sliced into 4-5 pieces and then placed into an inverted funnel submerged at the base in previously de-gassed deionized water (Supplementary File 9). Headspace air was removed through a septum in the top of the funnel. Subsequently, the vacuum chamber was sealed, and the aspirator used to extract the gas from within the fruit. 0.5mL headspace gas that collected in the neck of the inverted funnel was extracted in triplicate with 1mL hypodermic syringes.

Gas samples (2 replicates per treatment, 1/4 pear sections from 4 fruits per replicate) were injected into a HP 5890A gas chromatograph (Agilent, Avondale, PA, USA) with a flame ionization detector (FID) connected to a 0.53 mm × 15 m GS-Q-PLOT column (Agilent) to measure the ethylene composition of the extracted gas. Ethylene gas composition measurements were repeated every three days throughout each experimental time course. Mean and standard error for ethylene evolution were calculated for each sample time point.

### Firmness measurements

Firmness measurements were collected at each time point using four replicate pears from each treatment group. A GS-14 Fruit Texture Analyzer (GÜSS Instruments, South Africa) with an attached 8.0 mm probe set at 5.0 mm flesh penetration was used to measure firmness at 3 equidistant points around the equatorial region of each fruit following peel removal. Mean firmness value for each fruit was used for the final data assessment.

### °Brix

Soluble solid content was measured at each sample point by pooling approximately 0.5mL extracted juice from four replicate fruit per treatment/control and quantifying *°Brix* using a handheld refractometer.

### Statistical analysis of physiological data

Analysis of variance (ANOVA) was conducted for CO_2_, ethylene, firmness, and °Brix data within and across each of the experiments using SAS® University Edition (SAS Institute Inc., Cary, NC) with time, and GLA as treatments. Standard error for carbon dioxide evolution was plotted in 3-day intervals, although measurements of CO_2_ were taken every 4 hours throughout the experiment.

### RNA Extraction and Sequencing

Total RNA was extracted from pulverized ‘D’Anjou’ and ‘Bartlett’ peel tissue for each of the three technical replicates at ‘Unripe’, ‘50% Ripened’, and ‘100% Ripened’ stages following fruit tissue specific DEPC-CTAB protocol [59]. RNA was quality checked on an agarose gel and was quantified using Nanodrop 2000 spectrophotometer (Thermo Scientific, Waltham, MA, USA). Following quality validation and quantification using a Life Technologies Qubit Fluorometer (Carlsbad, CA) and Agilent Bioanalyzer (Santa Clara, CA), RNA libraries were sequenced by BGI Hong Kong Tech Solution NGS Lab on an Illumina HiSeq 4000 platform as 2×100 paired end reads.

### Transcriptome Assembly

The 2×100 paired end fastq files generated using Illumina HiSeq 4000 were input into the CLC Bio Genomics Workbench (ver 6.0.1) (Aarhus, Denmark) for pre-processing and assembly. The CLC Create Sequencing QC report tool was used to assess quality. The CLC Trim Sequence process was used to trim quality scores with a limit of 0.001, corresponding to a Phred value of 30. Ambiguous nucleotides were trimmed, and the 13 5’ terminal nucleotides removed. Reads below length 34 were discarded. Overlapping pairs were merged using the ‘Merge Overlapping Pairs’ tool, and a subsequent de novo assembly was performed with all datasets. Parameters used in the assembly are as follows: Map reads back to contigs = TRUE, Mismatch cost = 2, Insertion cost = 3, Deletion cost = 0.4, Similarity Fraction = 0.95, Global Alignment = TRUE, Minimum contig length = 200, Update contigs = TRUE, Auto-detect paired distances = TRUE, Create list of un-mapped reads = TRUE, Perform scaffolding = TRUE. The de novo assembly resulted in the production of 148,946 contiguous sequences (contigs). Contigs with less than 2x coverage and those less than 200bp in length were eliminated. For each individual dataset (treatment/replicate) the original, non-trimmed reads were mapped back to the master assembly subset. Default parameters were used, except for the length fraction, which was set to 0.5, and the similarity fraction, which was set to 0.9. Mapping resulted in the generation of individual treatment sample reads per contig. The master transcriptome was exported as a fasta file for functional annotation and the read counts for each dataset were exported for normalization with the Reads Per Kilobase per Million reads (RPKM) method [60].

### Functional annotation

The master transcriptome fasta produced from the Illumina assembly was imported into OmicsBox 1.1.135 (BioBam Bioinformatics S.L., Valencia, Spain) for functional annotation of expressed contigs. Contig sequences were identified by a blastx alignment against the NCBI ‘Viridiplantae’ database with and e-value specification of 10.0E-3. Gene ontology (GO) annotation was assigned using the ‘Mapping’ and ‘Annotation’ features using default parameters to generate a functionally annotated master assembly [61].

### Differential expression analysis

Temporally differentially expressed genes were identified using the time course, multi-series differential expression feature in the OmicsBox suite, which employs the maSigPro R package [62]. FDR cutoff value was set to 0.05. The statistical analysis ensured that genes that did not meet the assumption of equal variances were eliminated from the analysis, which was particularly important given that the three experiments were performed at different times throughout the 2018 season. The DEGs and expression values were matched with their corresponding functional annotations (Supplementary File 5).

### GO enrichment analysis

Gene ontology (GO) enrichment analysis was conducted to determine over and underrepresented biological processes, molecular functions, and cellular components among the differentially expressed sequences using the OmicsBox Enrichment Analysis (Fisher’s Exact Test) function [61] (Supplementary File 8). The annotated master transcriptome was used as the reference dataset, and the set of genes identified as differentially expressed over time in the treatment group versus the control group was used as the test dataset.

### qRT-PCR Validation

Primers for qRT-PCR targeting seven differentially expressed genes in the ripening-related pathways discussed previously were designed using the NCBI Primer-BLAST tool [63]. 200ng RNA for each sample was used to generate 1st strand cDNA using the Invitrogen VILO kit (Life Technologies, Carlsbad, CA USA). cDNA preparations were then diluted to 20ng/uL. Final library concentrations were quantified using a Qubit fluorometer (Carlsbad, CA). qRT-PCR technical replicate reactions were prepared for each of the gene targets using the iTAQ Universal SYBR Green Supermix with ROX reference dye (BioRad, Hercules, CA) per the manufacturer’s protocols with 20ng of template cDNA. In a Strategene MX3005P, the following thermocycle profile was used: 95°C initial disassociation for 2:30 minutes followed by 50 amplification cycles (95°C for 30s, 60°C for 30s, and 72°C for 30s) and a final, single cycle phase to generate a dissociation curve (95°C for 30s, 57°C for 30s, and 72°C for 30s). The LinRegPCR tool was used to calculate the Cq values for each reaction [64, 65] (Supplementary File 10). Cq values which were calculated from efficiency scores below 1.80 or above 2.20 were considered sufficiently low in confidence and were deemed unacceptable and were omitted from the analysis.

## Supporting information

S1_Brix2018

S2_Sugar_OrgAcid_HPLC

S3_Additional_RC_pH_Experiments

S4_Pear_GlyoxylicAcid_Assembly txt

S5_Annotated_RPKMs_Mean_Stderr

S6_AspCysMet

S7_ERFs

S8_EnrichedGOs_GLAvsC

S9_GLA_Humidification

S10_qRTPCR

## Acknowledgements

The authors thank Blue Bird Growers (Cashmere, WA) for their support and providing pears for conditioning experiments and to Scott Mattinson for assistance with the gas chromatography work. Work in the Dhingra lab was supported in part by Washington State University Agriculture Center Research Hatch Grant WNP00011 and grant funding from Pear Bureau NW to AD. SLH acknowledges the support received from ARCS Seattle Chapter and National Institutes of Health/National Institute of General Medical Sciences through an institutional training grant award T32-GM008336. The contents of this work are solely the responsibility of the authors and do not necessarily represent the official views of the NIGMS or NIH.

## SUPPLEMENTARY FILES

**Supplementary File 1.** Soluble solid content of 1-MCP treated ‘D’Anjou’ pears following treatment with 3% GLA in February and June 2018 experiments.

**Supplementary File 2.** Mean HPLC profiles for glucose, fructose, citric acid, and malic acid for 2018 1-MCP ‘D’Anjou’ pears treated with 3% GLA solution.

**Supplementary File 3.** Additional results from GLA experiments and pH experiments conducted in the 2017 and 2018 pear seasons.

**Supplementary File 4.** Master transcriptome assembly fasta file for control and 3% GLA treated 1-MCP ‘D’Anjou’ pear fruit.

**Supplementary File 5.** Mean RPKM values, standard error, and time course differential expression information for GLA-treated and control ‘D’Anjou’ pear fruit.

**Supplementary File 6.** Genes involved in metabolism of amino acids leading into precursors of ethylene biosynthesis that were differentially expressed in the 3% GLA-treated 1-MCP ‘D’Anjou’ pear fruit versus control.

**Supplementary File 7.** Ethylene response factor encoding genes that were differentially expressed in 3% GLA-treated 1-MCP ‘D’Anjou’ pear fruit versus control.

**Supplementary File 8.** Enriched gene ontologies for GLA-treated pear fruit versus control pear fruit during experimental time course.

**Supplementary File 9.** Images of GLA ultrasonic humidification system, internal ethylene gas extraction method, and CO_2_ evolution monitoring system.

**Supplementary File 10.** Quantitative RT-PCR results with LinRegPCR output and calculated expression values.

