## Supplementary material for "Glyoxylic acid overcomes 1-MCP-induced blockage of fruit ripening in *Pyrus communis L*. var. ‘D’Anjou’": S3_Additional_RC_pH_Experiments

Experiments using glyoxylic acid solutions of various concentrations and pHs were conducted during the 2017 and 2018 pear seasons (Supplementary Table 1).  Three experiments conducted in 2017 pear season allowed for identification of optimal glyoxylic acid treatment solution and sampling time course to elicit ripening responses. Based on the observations that glyoxylic acid at its native pH of 2.3 elicited the most prominent ripening response, resulting in significantly decreased firmness, increased internal ethylene production, and CO_2_ evolution over time in comparison with the control, 3% glyoxylic acid solutions at native pH and pH values titrated to 4 and 6 were selected as the treatment solutions to be tested in the 2018 experiments.

The 2017 experiments revealed that as the concentration of glyoxylic acid applied to fruit increased in the 0 to 3% range, more dramatic visual responses were observed, particularly related to the aesthetic quality and textural composition of the fruit. Increased incidence of peel tissue burning was observed with higher glyoxylic acid concentrations (Supplementary Figure 1). Fruit physiology was also altered as a result of ripening compound, with fruit treated with glyoxylic acid at native pH (2.3) displaying the most dramatic decrease in firmness, increase in internal ethylene and increase in CO_2_ evolution over time, in comparison with fruit treated with glyoxylic acid solutions titrated to neutral pH and the control solution (Supplementary Figure 2a-c) or with solutions of 1% and 2% glyoxylic acid (Supplementary Figure 3a-c).

To test whether acidity of glyoxylic acid solutions was responsible for induction of ripening in the fruit, and to observe the effects of neutrally shifted pH on ripening, several pH titrations were employed in a trial with 1% GLA solutions at native pH and pH 6, along with a control solution of D-isoascorbic acid. The final compound was chosen as an acidity control because it is the isomer of ascorbic acid that is not metabolized by plants, and therefore serves to represent the effects of acidic compound without affecting metabolism of the fruit in other ways (Kka et al., 2017). Treatment with D-isoascorbic acid yielded results similar to the those of the control, in which the fruit did not display a significant ripening response in comparison with either of the 1% native pH solution or the 1% pH 6 solution, both of which displayed decreases in firmness and increases in internal ethylene production during the ripening experiments (Supplementary Figure 4a-b). The results of the preliminary trials conducted in 2017 suggested that the acidic nature of the glyoxylic acid treatment solutions is, in part, responsible for accelerated ripening of the fruit, potentially via alteration of the capacity of enzymatic reactions to take place; however, application a non-metabolizable acidic compound alone (D-isoascorbic acid) was not sufficient for induction of ripening, although it did result in notably less superficial damage to the outer surface of the fruit.

2018 pH experiments, although conducted at the same time as the control and 3% RC experiments used for transcriptome sequencing, were not reported in the manuscript. Results of those studies are reported here in addition to those of the 2017 experiments (Supplementary Figures 5-7).

**Supplementary Table 3.1** Informational table detailing 1-MCP ‘D’Anjou’ pear RC experiment date ranges, treatments applied during each experiment, and the growers from which pears were obtained.

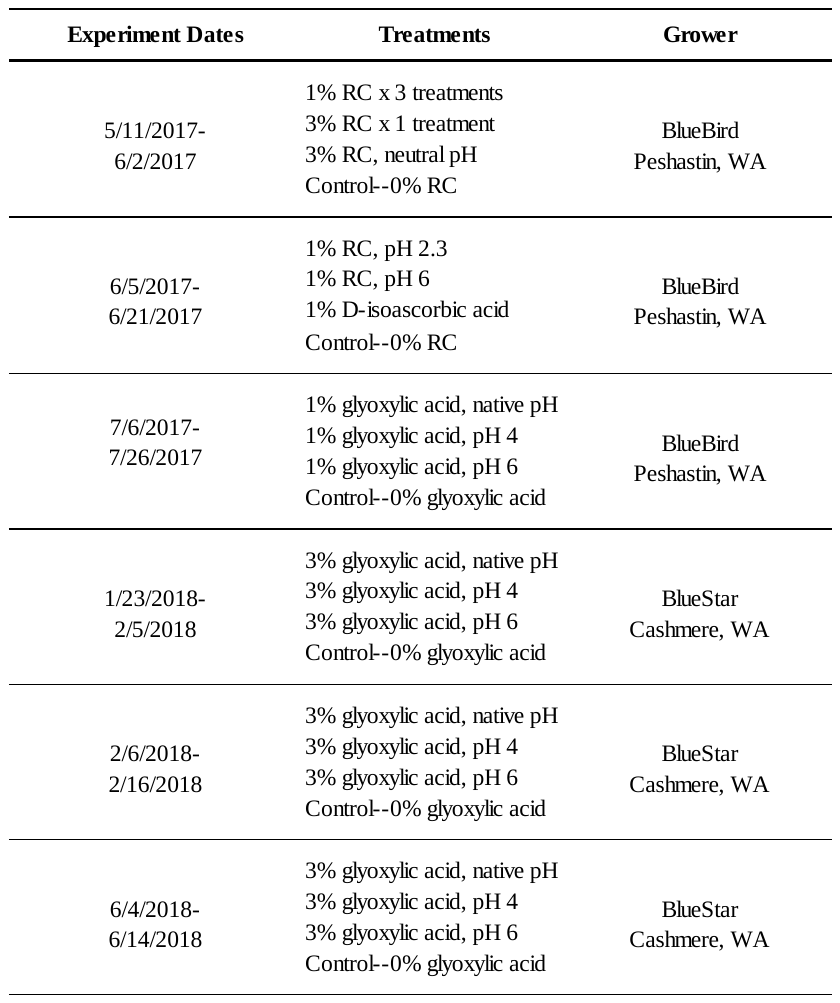

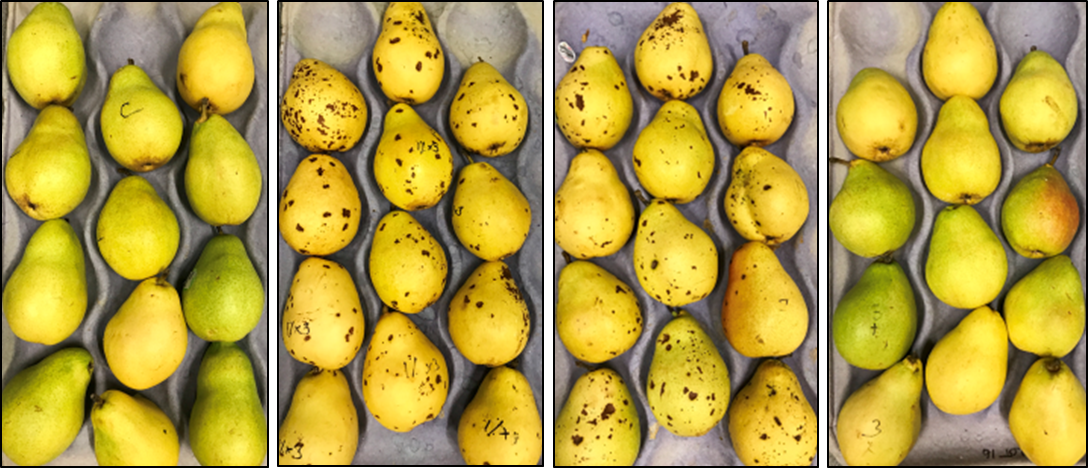

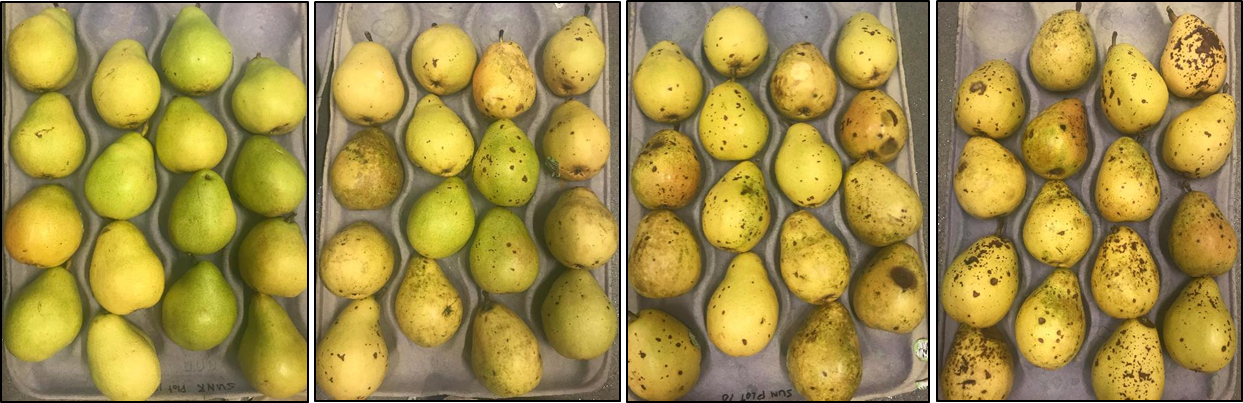

**Supplementary Figure 3.1.** Images taken on final day of a glyoxylic acid concentration experiment. Top— ‘D’Anjou’ pears at day 14 following glyoxylic acid/pH treatments. Left to right: control; 1% glyoxylic acid (three applications); 3% glyoxylic acid (native pH); 3% glyoxylic acid pH 6. Bottom— ‘D’Anjou’ pears at day 14 following glyoxylic acid treatments. Left to right: Control, 1% glyoxylic acid, 2% glyoxylic acid, 3% glyoxylic acid.

Firmness (lbf)

| Days After Treatment | Control | 1% x 3 treatments | 3% | 3%, neutral pH |
| --- | --- | --- | --- | --- |
| 1 | 14.0867 | 13.5 | 14.3258 | 13.79 |
| 20 | 9.00333 | 1.835 | 2.62125 | 7.94917 |

**Supplementary Figure 3.2a.** Starting and ending firmness of 1-MCP treated ‘D’Anjou’ pear fruit subjected to glyoxylic acid treatment in May 2017. Asterisks indicate significant difference from the control (p<0.05).

*****

*****

Firmness (lbf)

| Days After Treatment | Control | 1% x 3 treatments | 3% | 3%, neutral pH |
| --- | --- | --- | --- | --- |
| 0 | 0.362005 | 2.13643 | 0.7752 | 0.32348 |
| 20 | 1.70659 | 24.895 | 25.1417 | 1.59357 |

**Supplementary Figure 3.2b.** Internal ethylene concentrations of 1-MCP treated ‘D’Anjou pear fruit subjected to glyoxylic acid treatments in May 2017. Asterisks indicate significant difference from the control (p<0.05).

CO2 Flow (mL kg-1 hr-1)

| Days After Treatment | Control | 1% RC x 3 | 3% RC | 3% RC Neutral pH |
| --- | --- | --- | --- | --- |
| 1 | 1108.4 | 1654.68 | 1874.19 | 1570.52 |
|  | 991.846 | 1509.46 | 1611.24 | 1335.15 |
|  | 936.581 | 1435.68 | 1469.18 | 1164.43 |
|  | 932.126 | 1436.23 | 1447.47 | 1130.09 |
|  | 908.288 | 1401.8 | 1417.95 | 1133.87 |
|  | 890.866 | 1381.19 | 1392 | 1061.66 |
| 2 | 879.648 | 1360.2 | 1378.58 | 1042.13 |
|  | 880.326 | 1353.48 | 1370.55 | 1039.33 |
|  | 872.403 | 1340.12 | 1358.96 | 1017.84 |
|  | 869.698 | 1335.31 | 1348.39 | 1003.6 |
|  | 866.961 | 1331.01 | 1345.53 | 1000.76 |
|  | 856.629 | 1313.36 | 1324.94 | 981.981 |
| 3 | 848.071 | 1296.62 | 1306.51 | 968.098 |
|  | 848.241 | 1293.76 | 1299.43 | 965.599 |
|  | 839.318 | 1280.42 | 1283.54 | 950.591 |
|  | 836.398 | 1275.72 | 1274.19 | 943.496 |
|  | 836.959 | 1272.08 | 1269.3 | 938.851 |
|  | 829.688 | 1260.92 | 1253.56 | 927.361 |
| 4 | 825.861 | 1249.91 | 1238.26 | 917.989 |
|  | 826.668 | 1245.54 | 1230.05 | 915.119 |
|  | 820.826 | 1234.25 | 1215.2 | 904.109 |
|  | 820.771 | 1230.37 | 1208.67 | 897.851 |
|  | 823.323 | 1230.85 | 1203.23 | 899.911 |
|  | 820.994 | 1225.89 | 1190.78 | 890.478 |
| 5 | 813.961 | 1209.82 | 1171.7 | 874.413 |
|  | 821.619 | 1216.77 | 1177.47 | 872.971 |
|  | 814.524 | 1200.53 | 1157.56 | 867.538 |
|  | 818.081 | 1202.16 | 1155.17 | 867.591 |
|  | 822.433 | 1203.86 | 1152.9 | 866.578 |
|  | 833.001 | 1206.83 | 1138.97 | 859.021 |
| 6 | 816.566 | 1191.62 | 1127.2 | 847.208 |
|  | 818.321 | 1189.95 | 1121.1 | 845.159 |
|  | 814.851 | 1183.72 | 1108.84 | 838.499 |
|  | 816.526 | 1184.08 | 1102.3 | 835.181 |
|  | 819.718 | 1186.75 | 1101.97 | 835.763 |
|  | 822.708 | 1190.8 | 1100.9 | 831.543 |
| 7 | 824.604 | 1194.22 | 1093.15 | 828.631 |
|  | 827.719 | 1197 | 1088.21 | 823.389 |
|  | 827.446 | 1198.72 | 1082.1 | 820.806 |
|  | 829.481 | 1204.01 | 1080.9 | 819.716 |
|  | 836.171 | 1210.35 | 1082.15 | 819.326 |
|  | 841.876 | 1227.68 | 1088.42 | 822.901 |
| 8 | 845.681 | 1237.88 | 1090.26 | 823.893 |
|  | 848.051 | 1239.32 | 1082.44 | 822.951 |
|  | 851.406 | 1254.52 | 1087.46 | 821.771 |
|  | 855.458 | 1265.68 | 1089.2 | 820.708 |
|  | 861.361 | 1272.72 | 1089.62 | 823.346 |
|  | 868.681 | 1295.6 | 1101.34 | 826.301 |
| 9 | 873.806 | 1311.64 | 1103.12 | 826.233 |
|  | 876.359 | 1313.85 | 1098.2 | 824.231 |
|  | 880.549 | 1337.36 | 1106.33 | 826.391 |
|  | 884.884 | 1352.64 | 1111.11 | 827.896 |
|  | 929.213 | 1396.86 | 1130.1 | 835.261 |
|  | 949.261 | 1488.94 | 1215.94 | 835.261 |
| 10 | 949.261 | 1488.94 | 1215.94 | 835.261 |
|  | 949.261 | 1488.94 | 1215.94 | 835.261 |
|  | 949.261 | 1488.94 | 1215.94 | 835.261 |
|  | 949.261 | 1488.94 | 1215.94 | 835.261 |
|  | 949.261 | 1632.3 | 1260.12 | 835.261 |
|  | 949.261 | 1663.91 | 1272.5 | 917.236 |
| 11 | 995.431 | 1643.02 | 1260.52 | 904.721 |
|  | 994.941 | 1642.83 | 1255.57 | 899.594 |
|  | 999.973 | 1680.04 | 1276.27 | 903.329 |
|  | 1006.87 | 1703.8 | 1288.82 | 908.823 |
|  | 1013.95 | 1712.82 | 1293.01 | 914.746 |
|  | 1025.5 | 1760.95 | 1322.37 | 923.203 |
| 12 | 1033.29 | 1788.21 | 1335.25 | 927.734 |
|  | 1035.94 | 1793.59 | 1336.7 | 926.504 |
|  | 1041.84 | 1834.16 | 1360.28 | 934.116 |
|  | 1048.39 | 1858.12 | 1372.81 | 939.528 |
|  | 1056.34 | 1868.97 | 1378 | 943.361 |
|  | 1188.27 | 2045.47 | 1535.49 | 1075.47 |
| 13 | 1210.57 | 2096.15 | 1577.07 | 1099.36 |
|  | 1087.06 | 1963.54 | 1439.77 | 1105.77 |
|  | 1231.53 | 2141.83 | 1606.94 | 1114.63 |
|  | 1239.06 | 2162.86 | 1621.42 | 1121.55 |
|  | 1247.14 | 2175.91 | 1632.37 | 1130.93 |
|  | 1266.65 | 2225.56 | 1672.44 | 1149.7 |
| 14 | 1274.99 | 2247.38 | 1687.53 | 1156.52 |
|  | 1274.06 | 2248.36 | 1686.11 | 1153.1 |
|  | 1280.18 | 2280.56 | 1713.45 | 1161.42 |
|  | 1284.77 | 2297.14 | 1729.72 | 1166.34 |
|  | 1289.28 | 2302.91 | 1728.62 | 1067.52 |
|  | 1156.75 | 2186.02 | 1616.88 | 1037.12 |
| 15 | 1161.49 | 2203.54 | 1631.19 | 1042.74 |
|  | 1170.25 | 2210.44 | 1634.79 | 1040.15 |
|  | 1169.78 | 2230.61 | 1658.87 | 1053.56 |
|  | 1176.59 | 2243.72 | 1675.41 | 1063.55 |
|  | 1180.65 | 2249.43 | 1677.51 | 1063.51 |
|  | 1191.81 | 2275.08 | 1713.79 | 1077.47 |
| 16 | 1196.88 | 2287.5 | 1728.96 | 1085.03 |
|  | 1199.02 | 2289.43 | 1732.69 | 1083.34 |
|  | 1203.13 | 2303.89 | 1755.87 | 1094.49 |
|  | 1208.03 | 2312.29 | 1769.74 | 1101.59 |
|  | 1214.16 | 2313.06 | 1780.06 | 1108.01 |
|  | 1223.45 | 2336.77 | 1808.24 | 1125.04 |
| 17 | 1230.31 | 2348.79 | 1825.04 | 1133.66 |
|  | 1230.94 | 2348.53 | 1825.98 | 1134.47 |
|  | 1239.65 | 2363.17 | 1855.29 | 1147.74 |
|  | 1244.96 | 2370.92 | 1869.38 | 1156.44 |
|  | 1252.85 | 2379.37 | 1874.82 | 1162.98 |
|  | 1267.79 | 2397.53 | 1913.22 | 1179.94 |
| 18 | 1275.01 | 2407.14 | 1929.5 | 1190.49 |
|  | 1278.88 | 2411.16 | 1934.18 | 1188.13 |
|  | 1285.06 | 2417.59 | 1955.46 | 1202.76 |
|  | 1291.31 | 2420.42 | 1968.28 | 1211.68 |
|  | 1299.6 | 2425.13 | 1979.87 | 1218.88 |
|  | 1315.4 | 2442.73 | 2010.66 | 1242.33 |
| 19 | 1324.77 | 2448.92 | 2028.55 | 1251.05 |
|  | 1327.49 | 2448.82 | 2032.21 | 1249.82 |
|  | 1337.26 | 2455.27 | 2053.78 | 1266.56 |
|  | 1346.52 | 2455.88 | 2071.86 | 1277.61 |

**Supplementary Figure 3.2c** Carbon dioxide evolution of 1-MCP treated ‘D’Anjou’ pears over the course of 19 days in May 2017. (Day 20 is not shown, because not all replicates of each treatment were measured before the experiment was terminated). Error bars represent standard error measurements calculated for days 0, 4, 7, 10, 13, and 16 following treatment with glyoxylic acid ripening compound or control solutions. p<0.05 when comparing the 3% RC and 1% RC x3 treatments to the Control and 3% RC titrated to neutral pH.

Firmness (lbf)

| Days After Treatment | Control | 1% | 2% | 3% |
| --- | --- | --- | --- | --- |
| 1 | 13.4283 | 13.7708 | 13.2217 | 12.3508 |
| 16 | 11.9077 | 1.87208 | 3.03208 | 3.3325 |

**Supplementary Figure 3.3a.** Starting and ending firmness of 1-MCP treated ‘D’Anjou’ pear fruit subjected to glyoxylic acid treatments in June 2017.

Internal Ethylene (ppm)

| Days After Treatment | Control | 1% | 2% | 3% |
| --- | --- | --- | --- | --- |
| 1 | 0.01131 | 6.66452 | 2.73887 | 8.40105 |
| 4 | 0.44685 | 8.58622 | 3.72344 | 3.62246 |
| 7 | 0.184605 | 83.6465 | 13.1852 | 25.642 |
| 10 | 0.143338 | 61.688 | 39.2206 | 47.284 |
| 14 | 1.25871 | 21.472 | 34.337 | 36.7839 |

**Supplementary Figure 3.3b.**  Internal ethylene evolution of 1-MCP treated ‘D’Anjou’ pear fruit subjected to glyoxylic acid treatment in June 2017.

CO2 Flow (mL kg-1 hr-1)

| Days After Treatment | Control | 1% | 2% | 3% |
| --- | --- | --- | --- | --- |
| 1 | 1255.66 | 1821.34 | 1848.58 | 1956.85 |
|  | 1012.27 | 1548.98 | 1830.7 | 1959 |
|  | 896.931 | 1340.35 | 1435.06 | 1602.83 |
|  | 875.873 | 1319.33 | 1426.66 | 1597.58 |
|  | 888.964 | 1327.28 | 1448.34 | 1618.93 |
| 2 | 908.253 | 1367.24 | 1504.23 | 1673.24 |
|  | 908.783 | 1364.05 | 1511.5 | 1682.71 |
|  | 911.874 | 1367.11 | 1443.99 | 1614.09 |
|  | 895.306 | 1360.4 | 1512.3 | 1684.28 |
|  | 882.516 | 1351.99 | 1508.72 | 1679.32 |
|  | 862.921 | 1330.5 | 1563.23 | 1728.15 |
| 3 | 868.346 | 1360.53 | 1544.82 | 1710 |
|  | 858.738 | 1355.07 | 1547.65 | 1712.14 |
|  | 847.528 | 1367.55 | 1539.46 | 1509.97 |
|  | 840.791 | 1352.38 | 1557.05 | 1720.01 |
|  | 832.673 | 1357.63 | 1567.37 | 1716.51 |
|  | 808.176 | 1626.49 | 1519.84 | 1672.99 |
| 4 | 819.721 | 1396.79 | 1592.85 | 1757.61 |
|  | 814.326 | 1389.36 | 1598.26 | 1761.43 |
|  | 811.156 | 1387.99 | 1595.24 | 1752.8 |
|  | 800.753 | 1384.68 | 1608 | 1768.88 |
|  | 797.019 | 1390.11 | 1618.35 | 1779.77 |
|  | 795.816 | 1397.47 | 1619.83 | 1778.24 |
| 5 | 787.991 | 1402.64 | 1636.54 | 1797.44 |
|  | 779.254 | 1400.1 | 1638.69 | 1799.53 |
|  | 778.019 | 1406.35 | 1637.97 | 1795.01 |
|  | 768.183 | 1406.64 | 1652.02 | 1809.62 |
|  | 764.154 | 1408.46 | 1684.53 | 1851.01 |
|  | 762.811 | 1414.89 | 1660.29 | 1819.59 |
| 6 | 759.958 | 1427.68 | 1688.86 | 1847.38 |
|  | 757.418 | 1435.28 | 1700.01 | 1856.14 |
|  | 756.351 | 1440.09 | 1698.33 | 1849.96 |
|  | 754.831 | 1449.24 | 1724.07 | 1878.17 |
|  | 753.384 | 1450.52 | 1734.91 | 1890.39 |
|  | 752.276 | 1462.98 | 1729.9 | 1885.58 |
| 7 | 753.143 | 1476.39 | 1757.64 | 1910.47 |
|  | 751.184 | 1479.86 | 1767.38 | 1919.27 |
|  | 747.733 | 1481.25 | 1760.95 | 1910.91 |
|  | 748.364 | 1488.49 | 1782.8 | 1927.84 |
|  | 750.184 | 1502.19 | 1794.29 | 1939.39 |
|  | 750.936 | 1509.82 | 1798.21 | 1941.33 |
| 8 | 750.836 | 1519.48 | 1814.25 | 1952.4 |
|  | 750.261 | 1525.19 | 1821.5 | 1955.05 |
|  | 748.026 | 1526.62 | 1818.36 | 1951.47 |
|  | 750.113 | 1535.72 | 1832.85 | 1960.8 |
|  | 753.704 | 1542.13 | 1836.08 | 1967.89 |
|  | 754.206 | 1551.77 | 1844.37 | 1975.67 |
| 9 | 759.221 | 1567.91 | 1862.05 | 1982.8 |
|  | 760.149 | 1574.27 | 1870.39 | 1984.46 |
|  | 759.951 | 1580.01 | 1870.62 | 1987.9 |
|  | 765.036 | 1589.12 | 1881.26 | 1992.96 |
|  | 764.721 | 1590.55 | 1880.88 | 1994.77 |
|  | 763.641 | 1599.15 | 1884.57 | 2003.07 |
| 10 | 775.193 | 1614.45 | 1900.24 | 2005.94 |
|  | 779.366 | 1619.79 | 1905.74 | 2010.79 |
|  | 774.728 | 1616.09 | 1898.56 | 2014.89 |
|  | 789.771 | 1633.87 | 1917.99 | 2019.68 |
|  | 796.111 | 1639.93 | 1921.51 | 2023.06 |
|  | 795.031 | 1641.28 | 1901.4 | 2115.61 |
| 11 | 857.999 | 1748.15 | 2052.83 | 2164.5 |
|  | 864.984 | 1758.54 | 2062.09 | 2170.23 |
|  | 832.296 | 1711.86 | 1978.59 | 2109.03 |
|  | 870.848 | 1767.71 | 2062.63 | 2171.16 |
|  | 877.256 | 1774.19 | 2064.63 | 2172.62 |
|  | 882.066 | 1782.94 | 2075.77 | 2185.74 |
| 12 | 891.076 | 1794.01 | 2078.41 | 2185.89 |
|  | 897.376 | 1800.02 | 2084.13 | 2192.76 |
|  | 896.444 | 1798.97 | 2084.06 | 2198.17 |
|  | 905.701 | 1812.18 | 2089.92 | 2195.45 |
|  | 909.598 | 1817.91 | 2095.16 | 2196.57 |
|  | 912.051 | 1818.49 | 2099.69 | 2214.51 |
| 13 | 927.541 | 1840.89 | 2117.66 | 2218.11 |
|  | 935.159 | 1851.52 | 2128.01 | 2226.58 |
|  | 932.676 | 1847.77 | 2126.8 | 2236.57 |
|  | 947.956 | 1868.17 | 2144.25 | 2239.24 |
|  | 953.643 | 1875.58 | 2154.24 | 2246.17 |
|  | 956.823 | 1883.62 | 2162.25 | 2273.71 |
| 14 | 977.313 | 1905.22 | 2200.52 | 2290.36 |
|  | 985.856 | 1914.55 | 2211.95 | 2298.28 |
|  | 984.109 | 1896.43 | 2198.85 | 2305.24 |
|  | 1001.01 | 1930.61 | 2225.85 | 2309.93 |
|  | 1009.55 | 1938.82 | 2231.05 | 2315.14 |
|  | 1011.29 | 1934.58 | 2235.7 | 2327.41 |
| 15 | 1029.6 | 1958.97 | 2247.25 | 2333 |
|  | 1033.83 | 1963.76 | 2250.39 | 2334.62 |
|  | 1035.75 | 1958.37 | 2258.55 | 2330.89 |
|  | 1042.7 | 1968.62 |  | 2331.04 |

**Supplementary Figure 3.3c.** CO_2_ evolution (respiration) of 1-MCP treated ‘D’Anjou’ pear fruit subjected to glyoxylic acid treatments in June 2017.

Firmness (lbf)

| Days After Treatment | Control | D-isoascorbic acid | 1% pH 2.3 | 1% pH 6 |
| --- | --- | --- | --- | --- |
| 1 | 11.165 | 11.8033 | 12.3442 | 11.43 |
| 16 | 8.3425 | 9.1575 | 4.00083 | 3.58833 |

**Supplementary Figure 3.4a.** Firmness of 1-MCP treated ‘D’Anjou’ pear fruit subjected to glyoxylic acid treatments in July 2017. Asterisks indicate significant difference from the control (p<0.05).

Internal Ethylene (ppm)

| Days After Treatment | Control | 1% D-isoascorbic acid | 1% pH 2.3 | 1% pH 6 |
| --- | --- | --- | --- | --- |
| 1 | 0 | 1.20409 | 3.40961 | 0.88411 |
| 4 | 0.041715 | 0 | 0.551875 | 0.1641 |
| 11 | 0.168938 | 0 | 0.22269 | 0.61133 |
| 18 | 0.7482 | 0.20979 | 7.66675 | 17.616 |

**Supplementary Figure 3.4b** Internal ethylene concentration of 1-MCP treated ‘D’Anjou’ pears following treatment with glyoxylic acid in July 2017.

Firmness (lbf)

|  | Control | 3% | 3% pH 4 | 3% pH 6 |
| --- | --- | --- | --- | --- |
| Unripe Day 1 | 11.7675 | 11.2767 | 11.5883 | 11.2308 |
| 50% Ripened Day 7 | 8.0725 | 5.65 | 5.56667 | 7.46 |
| 100% Ripened Day 14 | 6.65 | 1.09083 | 3.72917 | 6.98417 |

**Supplementary Figure 3.5a.** Firmness of 1-MCP treated ‘D’Anjou pears following treatment with 3% glyoxylic acid at various pHs in January-February 2018.

Firmness (lbf)

|  | Control | 3% | 3% pH 4 | 3% pH 6 |
| --- | --- | --- | --- | --- |
| Unripe Day 1 | 12.14 | 12.0592 | 10.5558 | 12.725 |
| 50% Ripened Day 4 | 6.24083 | 5.6325 | 5.7775 | 3.61 |
| 100% Ripened Day 10 | 2.88167 | 1.02583 | 1.43083 | 2.47667 |

**Supplementary Figure 3.5b.** Firmness of 1-MCP treated ‘D’Anjou’ pears following treatment with 3% glyoxylic acid at various pHs in February 2018.

Firmness (lbf)

|  | Control | 3% | 3% pH 4 | 3% pH 6 |
| --- | --- | --- | --- | --- |
| Unripe Day 1 | 11.8817 | 13.07 | 11.4475 | 11.4373 |
| 50% Ripened Day 4 | 10.06 | 6.8925 | 7.95 | 7.75896 |
| 100% Ripened Day 7 | 7.50583 | 2.03583 | 2.96833 | 4.18146 |

**Supplementary Figure 3.5c.** Firmness of 1-MCP treated ‘D’Anjou’ pears following treatment with 3% glyoxylic acid at various pHs in June 2018.

Internal Ethylene (ppm)

|  | Control | 3% RC | 3% RC pH 4 | 3% RC pH 6 |
| --- | --- | --- | --- | --- |
| Unripe Day 1 | 1.20457 | 0.882855 | 0.69678 | 0.73074 |
| 50% Ripened Day 7 | 0.576765 | 1.38459 | 0.96951 | 0.944415 |
| 100% Ripened Day 14 | 0.61239 | 15.676 | 6.39861 | 2.01859 |

**Supplementary Figure 3.6a.** Internal ethylene of 1-MCP treated ‘D’Anjou’ pears following treatment with 3% glyoxylic acid at various pHs in January-February 2018.

Internal Ethylene (ppm)

|  | Control | 3% | 3% pH 4 | 3% pH 6 |
| --- | --- | --- | --- | --- |
| Unripe Day 1 | 0.0259525 | 2.85512 | 0.72811 | 2.36023 |
| 50% Ripened Day 4 | 0.397473 | 1.19533 | 0.102078 | 0.407808 |
| 100% Ripened Day 10 | 0.554088 | 34.1496 | 12.7789 | 0.842665 |

**Supplementary Figure 3.6b.** Internal ethylene evolution of 1-MCP treated ‘D’Anjou’ pears following treatment with 3% glyoxylic acid at various pHs in February 2018.

Internal Ethylene (ppm)

|  | Control | 3% | 3% pH 4 | 3% pH 6 |
| --- | --- | --- | --- | --- |
| Unripe Day 1 | 2.24676 | 6.18159 | 7.15941 | 1.14912 |
| 50% Ripened Day 4 | 2.3817 | 5.90164 | 1.62331 | 1.11591 |
| 100% Ripened Day 7 | 1.26261 | 10.6557 | 9.39747 | 7.62086 |

**Supplementary Figure 3.6c** Internal ethylene evolution of 1-MCP treated ‘D’Anjou’ pears following treatment with 3% glyoxylic acid at various pHs in June 2018.

ºBrix

|  | Control | 3% RC | 3% RC pH4 | 3% RC pH 6 |
| --- | --- | --- | --- | --- |
| Unripe Day 1 | 12.2 | 12.7 | 13 | 12.125 |
| 50% Ripened Day 4 | 11.475 | 12.825 | 13.6 | 13.55 |
| 100% Ripened Day 10 | 13.275 | 14.05 | 13.55 | 12.925 |

**Supplementary Figure 3.7a.** Soluble solid content of 1-MCP treated ‘D’Anjou’ pears following treatment with 3% glyoxylic acid at various pHs in February 2018.

°Brix

|  | Control | 3% | 3% pH 4 | 3% pH 6 |
| --- | --- | --- | --- | --- |
| Unripe Day 1 | 11.5 | 12.6 | 12.45 | 11.65 |
| 50% Ripened Day 4 | 11.9 | 12.75 | 13.075 | 11.85 |
| 100% Ripened Day 7 | 12.4 | 13.35 | 13.625 | 12.8 |

**Supplementary Figure 3.7b.** Soluble solid content of 1-MCP treated ‘D’Anjou’ pears following treatment with 3% glyoxylic acid at various pHs in June 2018.
