## Supplementary material for "Glyoxylic acid overcomes 1-MCP-induced blockage of fruit ripening in *Pyrus communis L*. var. ‘D’Anjou’": S9_GLA_Humidification

### Slide 1
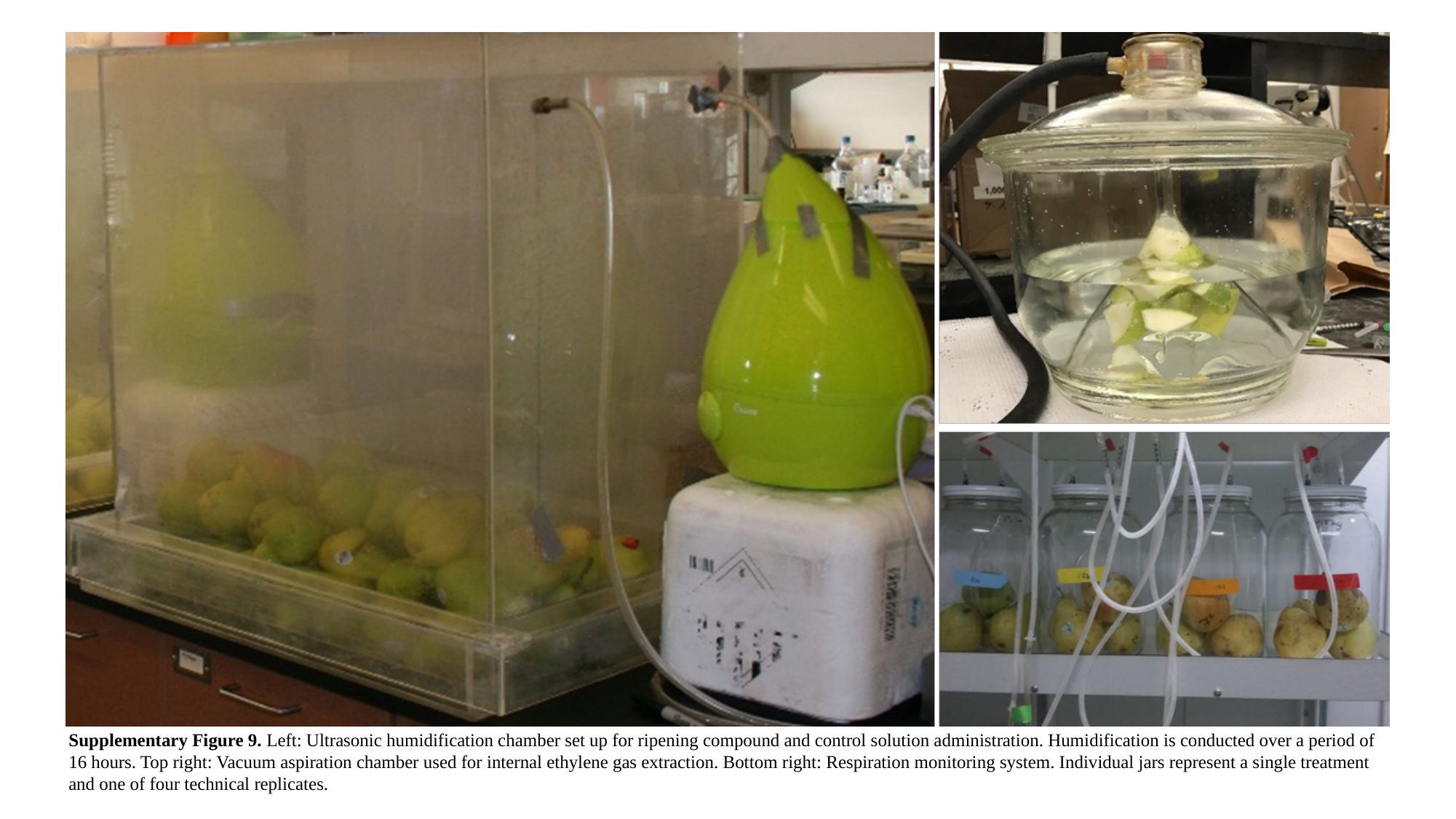

Supplementary Figure 9. Left: Ultrasonic humidification chamber set up for ripening compound and control solution administration. Humidification is conducted over a period of 16 hours. Top right: Vacuum aspiration chamber used for internal ethylene gas extraction. Bottom right: Respiration monitoring system. Individual jars represent a single treatment and one of four technical replicates.
